# In-cell structural insights into fungal ER stress responses

**DOI:** 10.64898/2026.04.10.717691

**Authors:** Leanne de Jager, Sofie van Dorst, Hannah Kugler, Marten L. Chaillet, Juliette Fedry, Stuart Howes, Friedrich Förster

## Abstract

Approximately a third of a cell’s proteome is folded in the endoplasmic reticulum (ER). The accumulation of misfolded proteins in the ER results in ER stress. The unfolded protein response (UPR) has evolved to attenuate ER stress by increasing the cell’s folding capacity and reducing the folding load in the ER. In metazoans, the latter is notably achieved through mRNA translation inhibition. However, *S. cerevisiae* lacks the corresponding pathways, and its reliance on translational downregulation in response to ER stress remains largely unexplored. In this study we combined cryo-focused ion beam milling, cryo-electron tomography and subtomogram averaging to investigate the effect of ER stress on the translational machinery of *S. cerevisiae*. At the ultrastructural level, we describe changes in organelle morphology, confirming ER volume expansion and the activation of autophagy. Further structural analysis reveals that ER stress triggers an increase in the relative abundance of inactive 80S ribosomes (up to 25% of the total), both in the cytosol and at the ER. Finally, we visualise the binding of the elongation factors eEF2, eIF5A and eEF3 to inactive ribosomes. Our work provides evidence that ER stress in *S. cerevisiae* leads to modest translation inhibition, which might contribute to alleviating ER stress.

## Introduction

Approximately 30% of the cell’s proteome utilizes the secretory pathway, including secreted proteins, plasma membrane proteins, and proteins destined for organelle membranes and lumens. After import into the endoplasmic reticulum (ER), the freshly synthesized peptide chains undergo various folding, maturation, and assembly steps. ER-specific and cytosolic chaperones aid peptide folding, the oligosaccharyltransferase mediates N-glycosylation, and protein disulfide isomerases catalyse the formation of disulfide bonds (Gemmer and Förster, 2020; Helenius et al., 1992). Together, these factors allow for efficient and accurate protein folding in the ER.

Protein folding, however, is inherently error-prone and inevitably results in occasional misfolded proteins (Hebert and Molinari, 2007). Quality control pathways help retain misfolded, and thus non-functional, proteins in the ER where they are either refolded or exported back into the cytosol for degradation (Adams et al., 2019). The ER faces continuous challenges from both physiological and pathological conditions which can result in an overload of misfolded proteins (Chen et al., 2011). This overload, caused by for example destabilizing mutations, aberrant lipid compositions affecting ER membrane integrity, and environmental challenges such as temperature fluctuations or pathogens, triggers ER stress. If ER stress remains unresolved, it will eventually lead to cell death (Chen et al., 2011).

To cope with ER stress, a highly conserved response termed the unfolded protein response (UPR) is activated. In metazoans, the UPR consists of three interacting branches, initiated by the proteins Inositol Requiring Enzyme 1 alpha (Ire1α), PKR-like ER kinase (PERK), and Activating Transcription Factor 6 (ATF6) (Cox et al., 1993; Harding et al., 1999; Haze et al., 1999). The accumulation of misfolded proteins results in the oligomerization and activation of Ire1α (Ire1p in yeast) as an RNase. Together with the tRNA ligase RTCB (Trl1p in yeast), Ire1α splices *XBP1* mRNA (*HAC1* in yeast), which gets translated into the transcription factor XBP1 (Hac1p in yeast). This results in the upregulation of a wide range of UPR-target genes (Jurkin et al., 2014; Lee et al., 2003; Sidrauski et al., 1996; Sidrauski and Walter, 1997). Together these genes increase the cell’s ER protein folding capacity and reduce its protein folding load (Travers et al., 2000).

ER stress and the UPR greatly affect our health. They are not only involved in various diseases, but also in basic processes like cell cycle progression and plasma cell differentiation (Hetz et al., 2013; Rutkowski and Hegde, 2010). To study the cell’s response to ER stress, *S. cerevisiae* is often used as a model system. The yeast’s UPR only consists of the Ire1p branch, which is evolutionary conserved (Ire1α in humans). This simplicity makes it more straightforward to study the UPR and makes yeast a powerful tool to dissect the conserved principles of ER stress signalling.

In yeast, activation of Ire1p results in the upregulation of nearly 400 UPR-target genes (Travers et al., 2000). The UPR increases the cell’s protein folding capacity via the upregulation of genes involved in protein folding and degradation, including the ERAD pathway (Berner et al., 2018), lipid metabolism, and vacuolar protein sorting. Interestingly, reduction of protein folding load, the second strategy by which ER stress is resolved in metazoans, has not been explored well in *S. cerevisiae*. In metazoans, a decrease in the protein folding load is achieved by reduction of mRNA translation initiation through eIF2α phosphorylation by PERK, and the degradation of ER-targeted mRNA by Ire1α (termed: Regulated Ire1α Dependent Decay, RIDD) (Hollien and Weissman, 2006; Wu et al., 2014). Since yeast lacks RIDD and the PERK pathway, translational control upon ER stress is thought to play a minor role in the yeast UPR (Maurel et al., 2014; Wu et al., 2014).

It is plausible, however, that translational control in yeast can be achieved in alternative ways. UPR target genes might affect translation indirectly. For example, UPR activation results in moderate downregulation of ribosomal proteins, which affect translation to a certain extent (Platzek et al., 2025; Travers et al., 2000). Moreover, first evidence of ER-stress induced translational control in yeast was provided recently, in the form of increased ubiquitination of ribosomal proteins upon ER stress (Matsuki et al., 2020). However, the possible impact of ER stress on global translation dynamics remains largely unknown. In this study we used cryo-focused ion beam (cryo-FIB) milling and cryo-electron tomography (cryo-ET) to explore the effect of ER stress on an ultrastructural level, and on both cytosolic and ER-localized translation in yeast.

## Results and Discussion

### Chemical treatment triggers ER stress and the UPR in S. cerevisiae

To study the effect of ER stress on translation, we chemically induced ER stress by treating yeast cells with chemicals that induce protein misfolding in the ER, thereby activating the UPR (van Anken et al., 2014). Cells were treated either short term (45 min.) or long term (4 hrs) with the reducing agent dithiothreitol (DTT), which prevents the formation of disulfide bonds in the ER. Alternatively, cells were treated with the glycosylation inhibitor tunicamycin (Tm) (3 hrs), which interferes with the biogenesis of many glycoproteins in the ER.

The UPR was monitored based on two of its hallmarks. Firstly, we measured the levels of spliced *HAC1* mRNA with RT-PCR (Sidrauski and Walter, 1997; Xia, 2019). Secondly, we used fluorescence microscopy (FM) to monitor the formation of Ire1p puncta resulting from ER stress (Aragon et al., 2009). To this end we performed most of our experiments (DTT, 45 min. and 4 hrs) using a yeast strain with C-terminally GFP-tagged Ire1p (Ire1c-GFP) (Huh et al., 2003). To confirm that our observations are not restricted to this specific strain, we also performed DTT treatment on a different strain (Ire1i-GFP) (Aragon et al., 2009; Halbleib et al., 2017), which shows higher *HAC1* splicing efficiency (Figure 1A, Supp. Fig. 1B). For Tm treatment, we generated a third strain using the brighter fluorophore NeonGreen (Ire1i-NG) (Supp. Fig. 1A). For all treatments and strains, we verified efficient *HAC1* mRNA splicing (Figure 1A, Supp. Fig. 1B-C) and the formation of Ire1p fluorescent punctae (Supp. Fig. 1E-I). Therefore, all conditions were included in our study.

**Figure 1:**
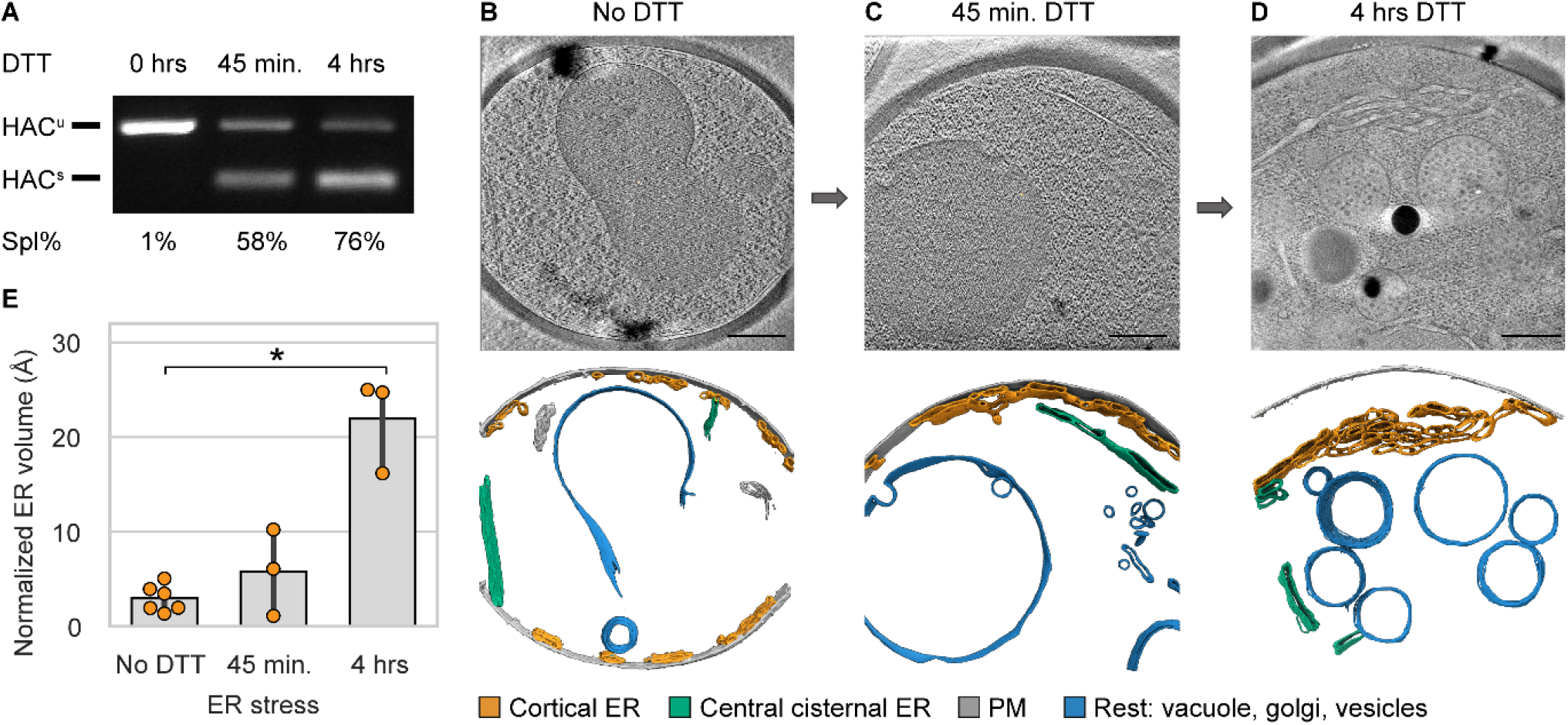
DTT treatment of yeast (Ire1c-GFP) induces the UPR. **(A)** RT-PCR of HAC1 mRNA isolated from untreated Ire1_c_-GFP cells and cells treated with 10 mM DTT for 45 min. or 4 hrs. Non-spliced HAC1 (HAC^u^, 433 bp) and spliced HAC1 (HAC^s^, 188 bp) can be detected. Splicing percentage is given below the image. **(B-D)** Slice through tomograms and corresponding organelle membrane segmentations of **(B)** a non-stressed cell, **(C)** a cell after 45 min. and **(D)** 4 hrs of DTT treatment. Segmented organelles are coloured: cortical ER (orange), central cisternal ER (green), plasma membrane (PM, grey) and remaining organelles (blue). **(E)** Quantification of normalised ER volumes in segmentations from non-stressed cells (N=6), and cells treated with DTT for 45 min. (N=3) or 4 hrs (N=3), (*P value <0.05, Mann Whitney U-test). Scalebar: 500 nm (B-D).

### *In situ* cryo-ET of *S. cerevisiae* confirms ER expansion and autophagy activation upon ER stress

To study the impact of the UPR at high resolution, we imaged treated and untreated yeast cells with cryo-ET. For DTT-treated Ire1c-GFP cells, tomograms were initially collected at relatively low magnification, capturing a near-complete cellular cross section. This strategy allowed us to study the effect of ER stress on cell morphology over time.

To cope with the increased ER folding load, the UPR induces major morphological changes at the ER, including an expansion of the ER volume (Bernales et al., 2006; Travers et al., 2000). *In situ* cryo-ET preserves the organelle membranes in a native hydrated state, making it suitable for the morphometric analysis of organelles. To visualise and quantify the impact of DTT stress on ER volume and morphology, untreated and treated Ire1c-GFP yeast cells were cryo-fixed at log phase by plunge freezing, prior to thinning by cryo-FIB milling. Cells were then imaged with cryo-ET at an intermediate magnification (6.32-7.09 Å/pix, FOV: ∼9 µm2), allowing us to laterally capture near-complete cellular ultrastructure in each tomogram (Figure 1B-D).

In our UPR-activating stress conditions, we found that the cortical ER expands considerably (Figure 1E). We observed a ∼7.4-fold increase after 4 hrs of ER stress, associated with the formation of parallel sheets (Figure 1D). No significant expansion was seen after 45 min., likely because the increase in lipid biosynthesis first requires a transcriptional change in lipid synthesis genes which only then leads to an increase in lipid abundance (Cox et al., 1993). The increase in ER volume upon 4 hrs of ER stress is in line with previous observations in chemically fixed cells by conventional EM, which report a 5-fold increase in ER volume after 3 hrs of DTT treatment (Bernales et al., 2006; Schuck et al., 2009).

In yeast, ER stress also triggers autophagy, which leads to the degradation and recycling of cellular components such as protein aggregates, and damaged or superfluous ER in the vacuole (Bernales et al., 2007; Høyer-Hansen and Jäättelä, 2007; Yorimitsu et al., 2006). Upon inspection of the vacuoles in our tomograms, we noticed distinct vacuolar morphological changes after ER stress, such as membrane invaginations and highly variable textures (Figure 1C-D, Supp. Fig. 4B and 7A). Moreover, we observed vacuoles that had taken up many ribosomes (both in the Ire1i-GFP strain after 4 hrs of DTT treatment, Supp. Fig. 7A-B, and the Ire1i-NG strain after 3 hrs of Tm treatment). These ribosomes were contained within autophagic bodies probably filled with cytosol. These observations are in line with the activation of autophagic responses upon ER stress in yeast, and suggest that autophagy may contribute to the active recycling of the translation machinery in these conditions.

### *In situ* subtomogram averaging visualises the native translation elongation cycle in *S. cerevisiae*

To investigate the effect of ER stress on translation, tomograms were collected at higher magnification (2.17 Å/pix, FOV: ∼1 µm^2^). For all conditions, 80S ribosomal particles (Figure 2A-B) were localized using template matching, yielding a total of 127,344 particles across all samples, which resulted in a 7.1-Å resolution subtomogram average (Supp. Fig. 2A). Ribosomal elongation states were visualized using a subtomogram alignment and classification strategy previously used for human ribosomes (Fedry et al., 2024; Gemmer et al., 2023) (Supp. Fig. 3).

**Figure 2:**
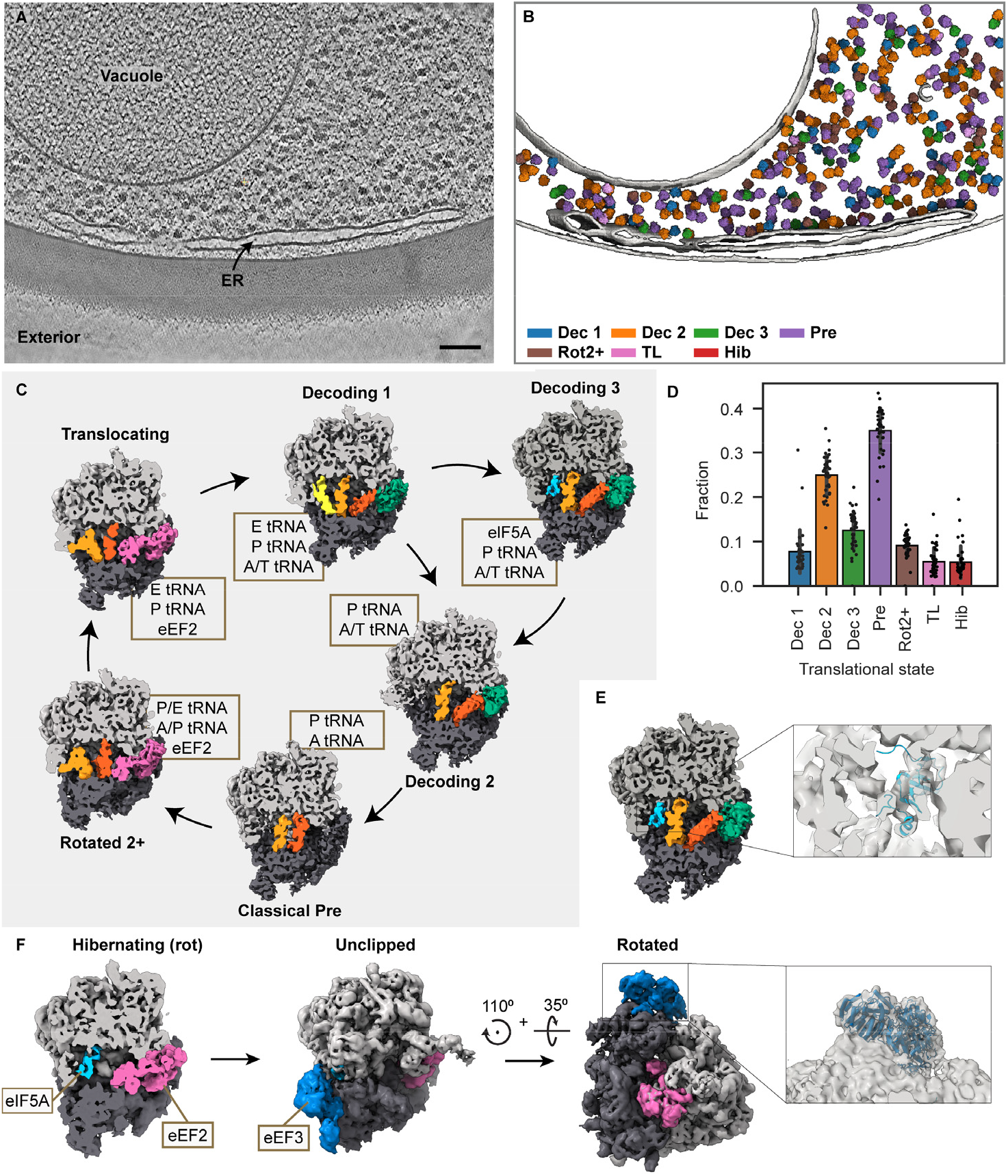
The yeast elongation cycle under non-stressed conditions. **(A)** Slice through a tomogram of the cytoplasm of a non-stressed cell (Ire1_c_-GFP). **(B)** Different ribosome elongation states mapped back in the tomogram shown in (A). Segmented membranes are displayed in grey. **(C)** The translational states found in our averaging workflow placed in a model elongation cycle. Ribosomes are clipped to visualise the tRNA sites. (**D**) The distribution of translational states found in non-stressed yeast cells is represented in a bar plot, individual data points are included (N=39 tomograms). **(E)** Zoom in on the Dec3 state with eIF5A (PDB: 5GAK) fitted into the density. **(F)** Clipped ribosome in a hibernating state. Unclipped view reveals an additional external density (dark blue). After rotating the ribosome 110^0^ around the Z-axis and 35^0^ around the X-axis, eEF3 (PDB: 7B7D, closed state) was fitted into this density. Scalebar: 100 nm (A-B).

Using this approach, we identified 7 distinct ribosome states (Figure 2C and F, Supp. Fig. 3A), corresponding to 6 elongating states and 1 inactive state. The active states capture the main steps of the elongation cycle, namely the decoding, peptidyl transfer and mRNA translocation steps. 6 out of the 7 states have been previously described *in vitro* and *in situ*, either in mammalian cells or in unicellular eukaryotes like *D. discoideum* and *S. cerevisiae* (Behrmann et al., 2015; Budkevich et al., 2014; Cheng et al., 2025; Fedry et al., 2024; Flis et al., 2018; Gemmer et al., 2023; Hoffmann et al., 2022; Rangan et al., 2024; Zheng et al., 2024), namely decoding (eEF1A, A/T, P tRNAs) state with or without E-site bound tRNA (respectively 7,255 and 23,809 particles in total) hereafter referred to as (i) *Dec1* and (ii) *Dec2*, (iii) classical PRE (*Pre* ; A, P tRNAs, 34,563 particles in total), (iv) rotated 2+ (*Rot2+* ; eEF2, A/P, P/E, 13,009 particles in total), (v) translocating (*TL* ; eEF2, P, E tRNAs, 5,389 particles in total) and (vi) hibernating (*Hib* ; eIF5A, eEF2, 18,463 particles in total). In addition to these known states, we observe a third decoding state named *Dec3* (18,652 particles) which was the third most abundant state in the non-stressed cells (Figure 2D). The *Dec3* state contains an A/T tRNA, a P-site tRNA, and an additional density close to the P-site tRNA (Figure 2E). This density is consistent in size, shape and location with eIF5A (local resolution of ∼7-8Å, Figure 2E, Supp. Fig. 2B) (Schmidt et al., 2015). eIF5A is a highly conserved translation elongation factor, which optimizes the iteraction between the P-site tRNA and the ribosome (Gregio et al., 2009; Henderson and Hershey, 2011). It increases the efficiency of translation, particularly upon ribosomes stalling at specific sequences like the poly-proline motif (Dever et al., 2014; Schuller et al., 2017). Hypusinated eIF5A is proposed to act by stabilising the P-site and favouring peptide bond formation, which is consistent with the previous *in situ* visualisation in *S. cerevisiae* of eIF5A in the classical PRE state (Rangan et al., 2024). Our abundant *Dec3* state suggests that eIF5A could also stabilise the P-site earlier in the elongation cycle, during the decoding step.

Overall, ∼95% of the ribosomes were classified in an elongating state in the non-stressed control cells (Figure 2D). This ratio is in good agreement with a recent cryo-ET study of translation elongation in *S. cerevisiae* (Cheng et al., 2025).

### An eEF3-, eIF5A- and eEF2-bound hibernating ribosome in *S. cerevisiae*

Inactive ribosomes are typically bound by several hibernation factors, protecting the ribosomes from degradation (Koli and Shetty, 2024). In yeast, Stm1 and Lso2 are well-characterised hibernation factors (Ben-Shem et al., 2011; Wells et al., 2020). They only bind the ribosome through small stretches of amino acids, and we could not confidently visualise such low molecular masses in our subtomogram averages of limited resolution. Nevertheless, our inactive 80S state, lacking bound tRNA’s, appears to be bound by the larger eIF5A and eEF2 (Figure 2F), which were previously observed at hibernating ribosomes in other unicellular as well as animal species, including humans (Du et al., 2024; Hoffmann et al., 2022; Leesch et al., 2023; Seraj et al., 2025).

Interestingly, we find an additional density enriched on hibernating rather than translating ribosomes. It was resolved at ∼9-15 Å resolution (Supp. Fig. 2C) and binds to the exterior of the ribosome, bridging the 40S and 60S subunit. Our assignment of this density as the eukaryotic elongation factor 3 (eEF3) is supported by a high confidence rigid body fit of a previous model of eEF3 bound to the yeast 80S ribosome (Figure 2F) (Ranjan et al., 2021). A recent *S. cerevisiae in situ* study visualised eEF3 bound to various elongating ribosomes (Cheng et al., 2025), consistent with its previously established role in translational regulation, including translocation, termination and recycling (Kobayashi et al., 2023; Kurata et al., 2010; Ranjan et al., 2021). The enrichment of eEF3 on inactive ribosomes in our data suggests that this factor may additionally play a role in yeast ribosomal hibernation.

### ER stress leads to a modest increase in global ribosome hibernation

To assess how ER stress affects translation over time, we next investigated how the distribution of translational states changes in the cytosol of yeast cells upon short-term (45 min. DTT, Figure 3A-B) and long-term (4 hrs DTT, Figure 3C-D) ER stress, compared to non-stressed cells. We observed an increase in the relative abundance of hibernating ribosomes (from 5% to 25%) at the expense of some of the major elongating states, namely *Dec2* and *Pre* (from 25% to 16% and from 35% to 20% (Fig. 3E, Supp. Fig. 4D). Similar changes in state abundances were also observed in Ire1_i_-GFP cells (45 min. and 4 hrs DTT, Supp. Fig. 4A-B) and upon 3 hrs Tm stress in Ire1_i_-NG cells (Supp. Fig. 4C). Collectively, these findings indicate that the observed translational response to ER stress is reproducible and generalizable across different conditions.

**Figure 3:**
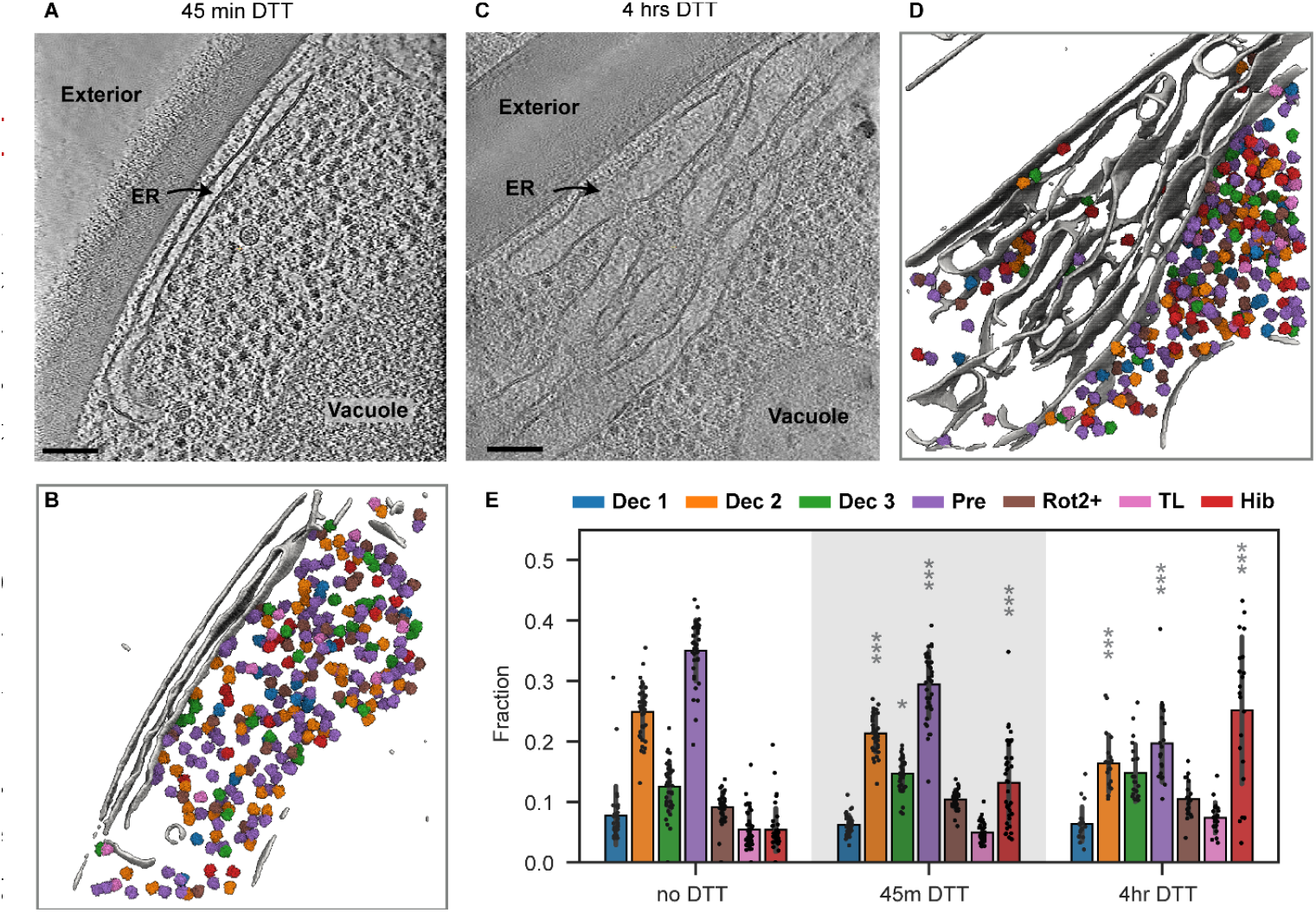
ER stress results in an increase in hibernating ribosomes in yeast cells. Slice through a tomogram of the cytoplasm of a yeast cell (Ire1_c_-GFP) stressed with DTT for 45 min. **(A)** or for 4 hrs **(C)**, including the segmented membranes and mapped-back ribosomes, coloured according to their translational state **(B, D)** see colour legend in (E). **(E)** The distribution of translational states found in different samples is represented as fractions in a bar plot (N=39, 37 and 21 tomograms for no DTT, 45 min. DTT and 4 hrs DTT respectively). States in stressed samples that had a significantly different abundance in comparison to the no DTT cGFP sample are indicated with grey asterixis (*P value <0.05, **P value <0.01, ***P value <0.001, Mann Whitney U-test). Scale bar: 100 nm (A-D).

In human cells, a similar DTT treatment was shown to result in near-complete hibernation of the cellular ribosomes (Gemmer et al., 2023). The lower hibernation rate in yeast hints at a more subtle mechanism of translational inhibition and a greater tolerance to the chemical stress compared to mammals.

Hibernating ribosomes are indicative of a reduced translation activity in the cell and can result from a block in translation initiation or ribosome recycling (Koli and Shetty, 2024). Here, initiation inhibition could occur through a general stress response (GSR), as the UPR interacts with other stress pathways in yeast (Pincus et al., 2014). The GSR can trigger Gcn2p-mediated phosphorylation of the initiation factor eIF2α (Cherkasova and Hinnebusch, 2003; Uppala et al., 2021). Additionally, upon long term ER stress the UPR or protein kinase A (PKA) might induce transcriptional changes of proteins involved in translational control, which can facilitate translational regulation (Pincus et al., 2014; Travers et al., 2000).

### ER stress similarly impacts ER-localised and cytosolic translation

As the UPR is aimed at specifically reducing ER stress, we next asked whether translation inhibition might be stronger at the ER membrane (Figure 4A-B). To address this question, we first aimed at visualising ER-bound ribosomes. We separated these from cytosolic ribosomes by focused image classification, using an ellipsoid mask positioned below the exit tunnel (see Methods, and Supp. Fig. 3C and 5A). Alignment and averaging of these particles yielded a ribosome with a membrane bilayer and an additional density at the exit tunnel corresponding to the ER translocon complex (Figure 4C, Supp. Fig. 2D, 6).

**Figure 4:**
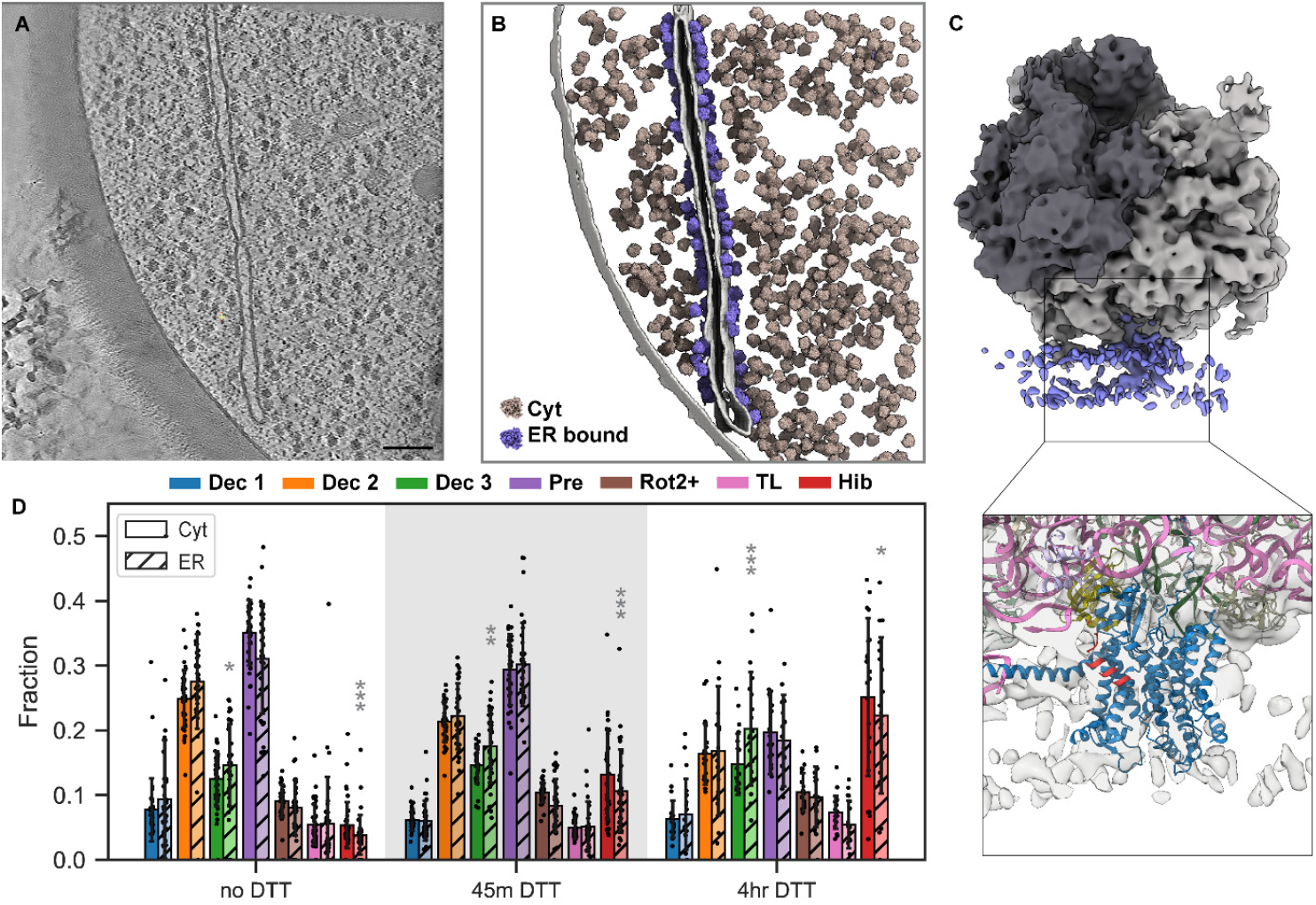
The impact of ER stress on ER-localised translation is similar to that on cytosolic translation. **(A)** Slice through a tomogram of a yeast cell (Ire1_c_-GFP) containing rough ER. **(B)** Corresponding membrane segmentation and mapped-back ribosomes coloured based on their assigned location, either cytosolic (brown) or ER-bound (purple). **(C)** ER-bound ribosome subtomogram average. Non-ribosomal densities are coloured in purple, SSU in dark grey and LSU in light grey. Black box zooms in on the density beneath the exit tunnel, with an atomic model for the yeast ribosome and translocon complex fitted into the density (PDB 3J77 and AlphaFold model, see M&M). **(D)** Bar graph representation of the abundance of each translational state per condition and location. Empty bars represent the cytosolic fraction and hatched bars represent the ER-bound fraction (N=35, 35 and 19 tomograms for no DTT, 45 min. DTT and 4 hrs DTT respectively). States where the ER-bound abundance is significantly different from the cytosolic abundance are indicated with grey asterixis (*P value <0.05, **P value <0.01, ***P value <0.001, Wilcoxon signed-rank test). Scale bar: 100 nm (A-B).

Rigid body fitting of a yeast translocon model (the yeast ribosome PDB 3J77, with an AlphaFold model of the yeast translocon) in our density highlights several ribosome-translocon interactions. Binding primarily occurs through the evolutionarily conserved loops 6/7 and 8/9 of Sec61p (hSec61α), as well as the N-terminal helix of Sss1p (hSec61γ) (Supp. Fig. 6C1), and possibly the C-terminus of Ysy6p (hRAMP4) (Supp. Fig. 6C2) (Becker et al., 2009; Lewis et al., 2024). Accessory translocon components such as TRAPα (Lewis et al., 2024) could not be resolved, which may be due to the variability of ribosome binding to the translocon complex. This ribosome-translocon density map indicates that our classification strategy successfully selects for ER-bound ribosomes and that ER stress did not prevent the association of 80S ribosomes to the ER translocon complex.

We next determined the translational state distribution of these ER-bound ribosome particles. Overall, we observed a consistent, stress-independent, increase of the *Dec3* state at the ER for all Ire1_c_-GFP conditions (Figure 4D). This increase might point to a role for eIF5A to support the elongation of nascent chains during import, helping to prevent stalling and translocon clogging. A similar increase in *Dec3* was seen for the other conditions, although not significant (Supp. Fig. 5D). Our data also indicated a slight, but consistent, decrease in hibernating ribosomes at the ER membrane compared to the cytosol (Figure 4D, Supp. Fig. 5D), presumably due to the higher affinity of ribosome nascent chain complexes for the ER translocon compared to inactive ribosomes.

Finally, like for cytosolic ribosome states, we observed that upon ER stress the abundance of hibernating states at the ER increased over time at the expense of other translating states (*Dec2* and *Pre*). Collectively our data show that during ER stress in yeast, a decrease in co-translational nascent chain import into the ER is achieved through a global inhibition of translation, instead of an ER-specific one. It is possible that additional mechanisms specifically inhibit ribosome-independent import at the ER translocon during ER stress.

### Vacuolar-residing ribosomes appear upon long-term ER stress

Besides distinct morphological changes in vacuolar structures upon ER stress, we also observed the presence of ribosomes in autophagic bodies after 4 hrs of DTT stress (Ire1_i_- GFP). Analysis of the translational state distribution of these vacuolar ribosomes showed that they are predominantly in a hibernating state (Supp. Fig. 7C-D, N=4 tomograms). Whether vacuoles selectively take up hibernating ribosomes or rapid conversion into the hibernating state occurs upon uptake is unclear. It seems more likely that random ribosomes are sequestered by autophagy in the cytosol and they transition to the hibernating state within the autophagic bodies. Our observations suggest that the vacuolar pathway is not only used for the degradation of protein aggregates or damaged ER during ER stress (Schuck et al., 2014; Travers et al., 2000), but potentially also contributes to the reduction in protein folding load by degradation of ribosomes.

## Conclusions and perspectives

In this study, we used cryo-FIB milling and cryo-ET to investigate the impact of ER stress on translation elongation in *S. cerevisiae*. It was previously thought that translational control plays a negligible role in yeast, as the UPR in this organism lacks the PERK and RIDD pathways (Wu et al., 2014). However, our data suggests that ER stress does have an impact on translation, albeit to a lesser extent than in higher eukaryotes (Gemmer et al., 2023).

We describe an increase in cytosolic hibernating ribosomes, to up to ∼25% of the total 80S ribosome population under prolonged DTT- or Tm-induced ER stress. This observation indicates that *S. cerevisiae* likely inhibits translation initiation to some degree during ER stress, possibly first due to the activation of general stress pathways via the kinases Gcn2p or PKA (Patil et al., 2004; Pincus et al., 2014) that could be further supported by the UPR transcriptional program during prolonged stress. Approximately 22% of the ER-bound ribosomes were also in the hibernating state during prolonged ER stress, confirming that the global translation reduction results in a corresponding decrease in cotranslational protein import into the ER.

The hibernating ribosome state observed in our data is bound by eIF5A. eIF5A is not a UPR-target gene (Travers et al., 2000), still its activity and localization can be regulated via a wide range of post translational modifications, including acetylation, phosphorylation, hypusination, ubiquitination and SUMOylation (Dever et al., 2014; Seoane et al., 2024; Shang et al., 2014). Furthermore, it responds to nutrient availability, proteotoxic stress and heat shock stress (Barba-Aliaga et al., 2020; Seoane et al., 2024). The various levels of regulation and its involvement in other stress responses make eIF5A a likely candidate for the regulation of translation under ER stress.

In summary this study exemplifies the use of *in situ* cryo-ET to provide valuable molecular insights into the effect of ER stress in *S. cerevisiae*, guiding novel hypothesis for further investigations in the field.

## Acknowledgements

We thank Professor Robert Ernst and Mike Renne (Saarland University) for providing the Sc-Ire1i-GFP strain and for insightful discussions regarding yeast strain generation. We thank Oliver Pajonk (Heidelberg University) for guidance in strain design and the generation of the Ire1i-NG strain. We thank Professor Sebastian Schuck (Heidelberg University) for his insightful feedback on the manuscript. We thank Mathieu Baltussen (Radboud University) for help with figure design. We thank Max Gemmer (Utrecht University), Gijs van der Schot (Utrecht University), Lena Thärichen (Utrecht University) and Dimitros Vismpas (Utrecht University) for helpful discussions regarding ribosome averaging strategies. Imaging was performed at the Utrecht University EM Centre with support from C. Schneijdenberg, J. Meeldijk, and M. Bergmeijer. The Netherlands Electron Microscopy Infrastructure (grant 84.034.014) helped support access to the Netherlands Center for Electron Nanoscopy with support from operator Willem Noteborn. This work was supported by the European Union (Imagine Consortium (GA#101094250)) and the Dutch Ministry of Education, Culture, and Science (OCW) (Gravitation Consortium “FLOW” (024.006.036)). This paper was typeset with the bioRxiv word template by @Chrelli: www.github.com/chrelli/bioRxiv-word-template.

## Author contributions

L. de Jager: Conceptualization, Data curation, Formal analysis, Investigation, Methodology, Project administration, Validation, Visualization, Writing – original draft, Writing - review & editing. Sofie van Dorst: Data curation, Formal analysis, Investigation, Methodology, Project administration. Hannah Kugler: Data curation, Formal analysis, Investigation, Methodology, Project administration. Marten Chaillet: Formal analysis, Methodology, Resources. Juliette Fedry: Methodology, Resources, Writing - review & editing. S.C. Howes: Conceptualization, Funding acquisition, Project administration, Supervision, Writing – review & editing. F. Förster: Conceptualization, Funding acquisition, Supervision, Writing - review & editing.

## Competing interest statement

The authors declare no competing interests.

## Materials and Methods

### List of reagents

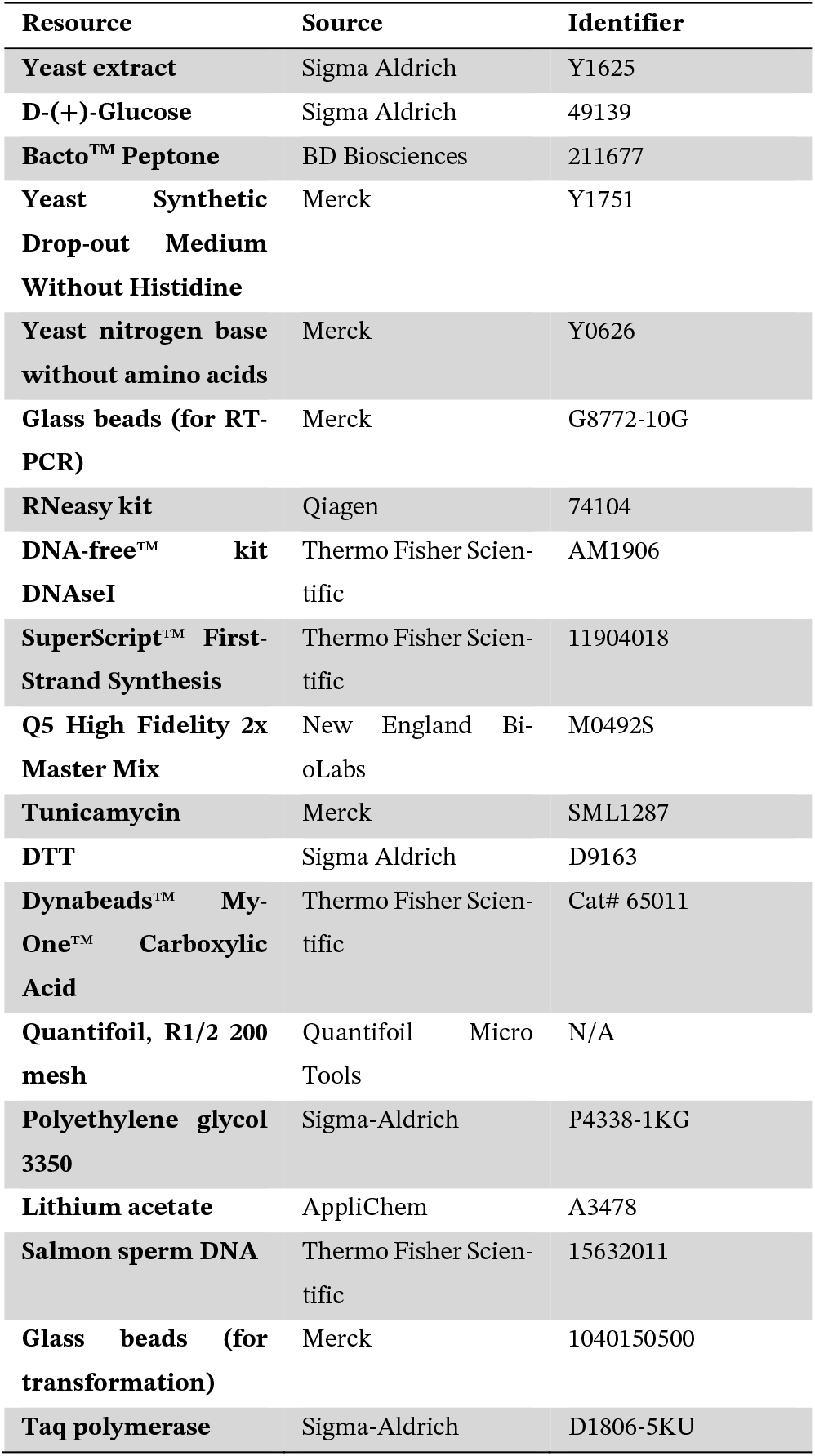

### Strains and cell culture

The *S. cerevisiae* strain BY4741 expressing Ire1p with a C-terminal GFP tag (Ire1C-GFP) was purchased from Thermo Fisher Scientific, originally part of the yeast GFP fusion localisation project (Huh et al., 2003). The *S. cerevisiae* strain BY4741 expressing Ire1p with an internal GFP tag (Ire1i-GFP) was a kind gift from Dr. Mike Renne and Prof. Robert Ernst (Saarland University) (Halbleib et al., 2017). The *S. cerevisiae* strain BY4741 expressing Ire1i-NeonGreen under control of the GAL1 promoter, located on a separate CEN/ARS plasmid, was generated in this study (Ire1i-NG). Ire1C-GFP and Ire1i-GFP strains were cultured in YPD medium ((1% yeast extract, 2% glucose, 2% peptone). The Ire1i-NG strain was grown in SC-His medium (2% glucose, 0.7% yeast nitrogen base, and amino acid mix without histidine).

### Plasmids and strain generation

To generate the plasmid pRS313-GAM1-IRE1_i_-NG-ADH1, the plasmid pRS313-IRE1_i_-NG was used as backbone and was PCR amplified using primer set 1 (Mathuranyanon et al., 2015). Using primer set 2, the GAM1 promoter was PCR amplified from plasmid pFA6a-kanMX6-GEM-PGAL1. Using primer set 3, IRE_i_-NG was PCR amplified from pRS313-IRE1_i_-NG. Using primer set 4, the ADH1 terminator was PCR amplified from plasmid pFA6a-kanMX6-GEM-PGAL1. The backbone and the three constructs were assembled together via Gibson Assembly. To generate the Ire1_i_-NG strain, pRS313-GAM1-IRE1_i_-NG-ADH1 and pNH605-ADH-GAL4-ER-M2 (pDEP151, supplied by David Pincus) were transformed using the LiAc method (Ito et al., 1983). A yeast culture was grown in YPD medium (1% yeast extract, 2% glucose, 2% peptone) overnight into saturation. The next morning, cells were back-diluted in 5 mL YPD in order to reach an OD of 1 within the next four hrs. After four hrs of growth, cells were spun down at 5,000 x g, washed with one volume of water and centrifuged again, this time at 10,000 x g. After the supernatant was discarded, transforming DNA was added to the pellet (10% of the PCR volume of transforming DNA). Subsequently, cells were resuspended in 360 µl transformation mix (33% (w/v) polyethylene glycol 3350, 100 mM lithium acetate, 0.28 µg/mL salmon sperm DNA, freshly boiled before use) and subjected to heat shock at 42°C for 40 min. After heat shock, cells were spun down and resuspended in 1 mL YPD medium. From the 1 mL culture in YPD, 100 µL cells were plated on selective plates using glass beads. After 2 days at 30°C, colonies were restreaked on selective plates and subjected to colony PCR to check for successful genetic modification.

Colony PCR was performed to check the successful genetic modification of yeast strains. Single colonies were picked from restreaked colonies. Single colonies were resuspended in the colony PCR mix (20 mM Tris-HCl, pH 8.8, 10 mM (NH_4_)_2_SO_4_, 10 mM KCl, 2.5 mM MgCl_2_). PCR was performed with standard conditions supplied by the manufacturer of the Taq polymerase (Sigma-Aldrich). Next, 8 µL of the PCR product was loaded on a 2x TAE + 1% agarose gel to pick out the clones which produced PCR products with the correct length.

### Cryo-ET sample preparation

Ire1_C_-GFP and Ire1_i_-GFP cells were incubated over night at 180 RPM and 30^0^C in YPD medium. In the morning, cells were diluted to an OD_600_ of ∼0.15 so at the moment of plunging, approximately 6 hrs later, the culture would have an OD_600_ of ∼0.6. Ire1_C_-GFP cells were either left to grow undisturbed (non-stressed) or treated with DTT (10 mM) for 45 min. or 4 hrs before plunging, to induce ER stress. Ire1_i_-GFP cells were treated with DTT 45 min. or 4 hrs before plunging as well. Ire1_i_-NG cells were grown at 180 RPM and 30^0^C during the day in SCD-His medium. At the end of the day, the cells were diluted to an OD_600_ of ∼0.001 so the next morning, approximately 16 hrs later, the cells would still be in the growth phase. The next morning, cells were diluted to an OD_600_ of ∼0.1 so the OD_600_ at the moment of plunging would be ∼0.7. 5 hrs prior to plunging, cells were treated with estradiol (100 nM) to induce Ire1_i_-NG expression. 3 hrs prior to plunging, cells were treated with tunicamycin (3 µg/mL).

To prepare cryo-EM grids, 45 µL yeast culture was mixed with 3 µL Dynabeads (1:16 dilution). Next, 3.25 µL yeast culture was incubated on glow discharged copper grids (Quantifoil, R1/2 200 mesh) for 30 seconds. The grids were vitrified in liquid ethane/propane (37%:63%) after manual blotting for 5 seconds. The grids were stored in liquid nitrogen throughout the subsequent experiments. In parallel, 2x 1.5 mL yeast culture was spun down (5 min., 2000 x g), the pellet was flesh frozen and stored at -80 ºC for later use in Hac1 splicing assays to confirm UPR activation.

### HAC1 splicing assay

Cell pellets stored in parallel to the plunging experiments were thawed and lysed by resuspending the cells in 700 μL RTL buffer and 500 µL glass beads, and vortexing for 10 min. at 4°C. RNA was purified using the RNeasy Mini Kit and treated with DNA-*free*™ kit DNAseI to remove DNA. cDNA was synthesized using the SuperScript™ First-Strand Synthesis System for RT-PCR with random hexamer primers. The cDNA of the HAC1 intron was amplified via PCR using the primers 5′-CCGTAGACAACAACAATTTG-3′ and 5′-CATGAAGTGATGAAGAAATC-3′. The PCR product was run on a gel to determine the presence of spliced *HAC1* mRNA. Splicing percentage was calculated by determining the average signal of each RNA band in ImageJ by drawing a ROI around the bands and calculating their plot profile. Background (Bg) signal was subtracted from each band before the final calculations.

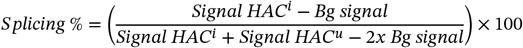

### Cryo-fluorescence microscopy

Grids were imaged with a cryo-fluorescence microscope (cryo-FM) either before milling (Ire1_c_-GFP 45 min.) or after the first 3 milling steps (rest of the samples). Cryo-FM data were collected before milling using a CorrSight confocal cryo-FM (FEI). This microscope is equipped with a cryo-stage, the Andromeda spinning disk module and a Hamamatsu ORCA-Flash4.0 camera. Z-stacks of intact grid squares containing large clusters of yeast were recorded. Ire1_c_-GFP and Dynabeads were excited at 488 nm. Data was collected using an EC Plan-Neofluar 40×/0.9 NA air objective, and the FEI MAPS v3.8 and LA FEI Live Acquisition v2.2.0 software (Thermo Fisher Scientific).

Cryo-FM data obtained after the first 3 milling steps was collected with the Meteor, a cryo-FM integrated in the cryo-FIB/SEM (Delmic). Z-stacks of the lamellae were recorded to detect Ire1p and the Dynabeads (LED excitation of 470 nm, emission filter 515/30). Data was collected using an Olympus LMPLFLN 50×/0.5 NA (Ire1_c_-GFP 4 hrs) or an Olympus LMPLFLN 50x/0.8 NA air objective (rest), and the Odemis 3.2.1 software (Delmic).

### Cryo-FIB Milling – manually or automated

Yeast cells were milled as established (Wagner et al., 2020); grids were first coated with a layer of platinum and organo-platinum. After this, milling was performed either manually or automated. Manually milled lamellae were produced in a stepwise manner by reducing the current from 1 to 0.3 to 0.1 nA and the thickness of the lamella from 2.5 to 1 to 0.5 µm per lamella site. Micro expansion joints were added during the first milling steps to increase lamella stability. Lamellae were polished at 30 pA to reach a thickness of 100-200 nm after which they were coated in platinum.

Automatically milled lamellae were produced using the AutoLamella python package (Buckley et al., 2020). In this case, micro expansion joints were made automatically first to improve the cross correlations of the software. After this, lamellae were produced in a stepwise fashion by reducing the current from 1 to 0.3 to 0.1 nA and the thickness of the lamella from 2.5 to 1.5 to 0.9 μm per lamella site. For automated milling details, see Table 3. In contrast to manual milling, where all milling steps at one lamella site were performed before moving to the next spot, the software performed the first milling step for all points of interest, then the second and then the third. The final polishing step, which reduced lamellae thickness to 100-200 nm, was performed manually at 30 pA. For the lamellae produced with AutoLamella, the first 5 min. of polishing were performed with the CERES ice shield (Delmic) inserted to reduce ice buildup. If lamellae stability permitted, polishing was continued for an additional 2 min. without the shield, guided by the SEM image of the lamella, after which they were coated in platinum.

**Table 1:**
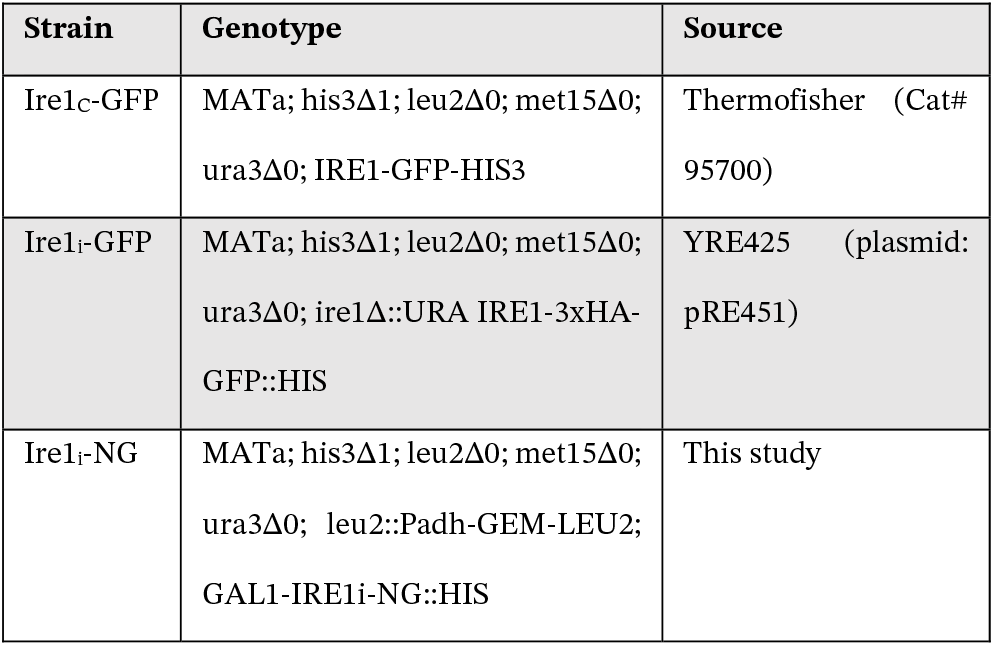
Strains used in this study.

| Strain | Genotype | Source |
| --- | --- | --- |
| Ire1C-GFP | MATa; his3Δ1; leu2Δ0; met15Δ0; ura3Δ0; IRE1-GFP-HIS3 | ThermoFisher (Cat# 95700) |
| Ire1i-GFP | MATa; his3Δ1; leu2Δ0; met15Δ0; ura3Δ0; ire1Δ::URA IRE1-3xHA-GFP::HIS | YRE425 (plasmid: pRE451) |
| Ire1i-NG | MATa; his3Δ1; leu2Δ0; met15Δ0; ura3Δ0; leu2::Padh-GEM-LEU2; GAL1-IRE1i-NG::HIS | This study |

**Table 2:**
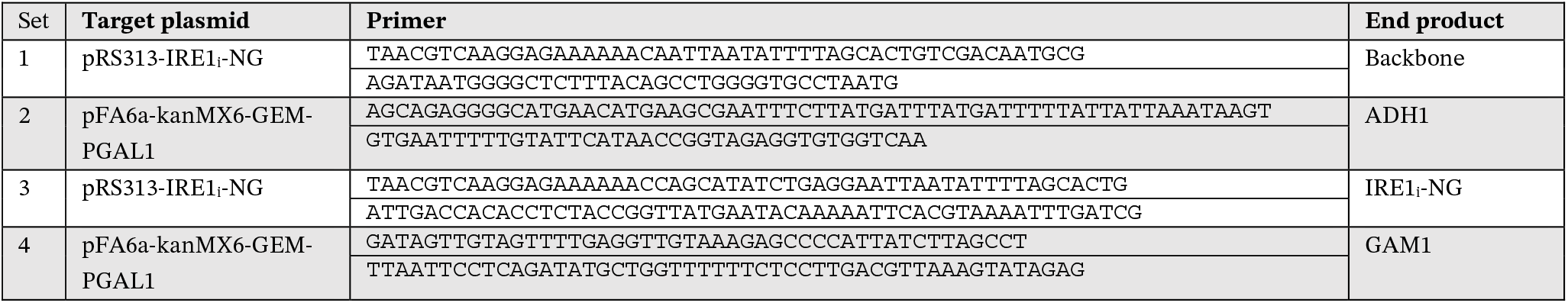
Primers used for cloning.

**Table 3:**
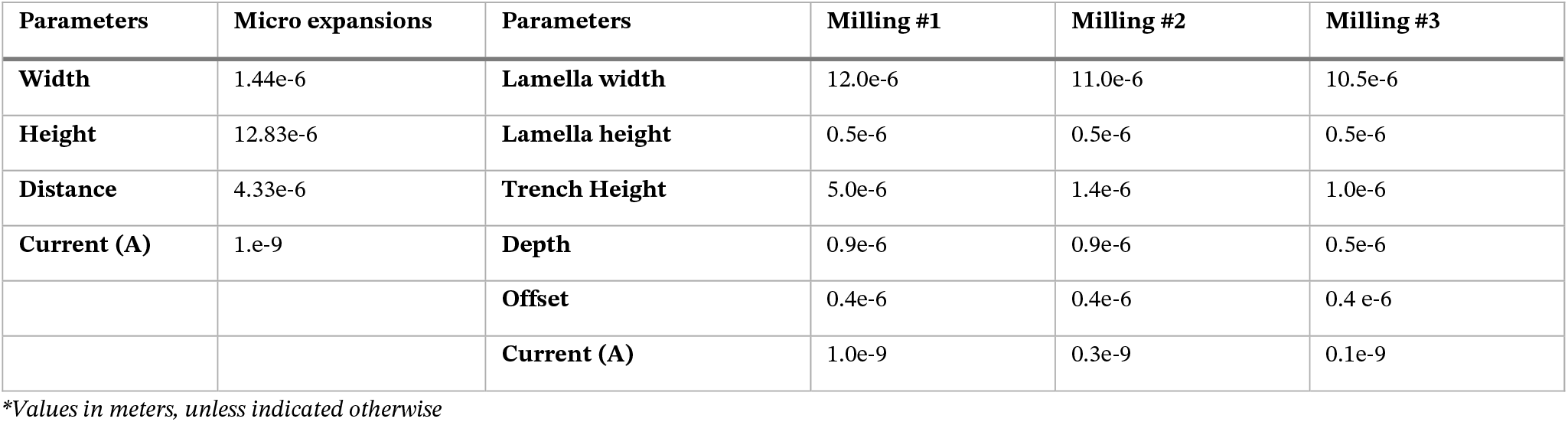
Milling parameters used with the AutoLamella package.

| Parameters | Micro expansions | Parameters | Milling #1 | Milling #2 | Milling #3 |
| --- | --- | --- | --- | --- | --- |
| Width | 1.44e-6 | Lamella width | 12.0e-6 | 11.0e-6 | 10.5e-6 |
| Height | 12.83e-6 | Lamella height | 0.5e-6 | 0.5e-6 | 0.5e-6 |
| Distance | 4.33e-6 | Trench Height | 5.0e-6 | 1.4e-6 | 1.0e-6 |
| Current (A) | 1.e-9 | Depth | 0.9e-6 | 0.9e-6 | 0.5e-6 |
|  |  | Offset | 0.4e-6 | 0.4e-6 | 0.4 e-6 |
|  |  | Current (A) | 1.0e-9 | 0.3e-9 | 0.1e-9 |

\*Values in meters, unless indicated otherwise

**Table 4:** Cryo-FIB and cryo-ET data collection parameters.

|  | *Ire1<sub>c</sub>-GFP*<br>No stress | *Ire1<sub>c</sub>-GFP*<br>45 min. DTT | *Ire1<sub>c</sub>-GFP*<br>4 hrs DTT | *Ire1<sub>i</sub>-GFP* 45 min.<br>DTT | *Ire1<sub>i</sub>-GFP*<br>4 hrs DTT | *Ire1<sub>i</sub>-NG*<br>3 hrs Tm |
| --- | --- | --- | --- | --- | --- | --- |
| Magnification | 63,000 or 42,000 | 63,000 or 42,000 | 63,000 or 42,000 | 42,000 | 42,000 | 42,000 |
| Voltage (kV) | 200 or 300 | 200 or 300 | 200 or 300 | 300 | 300 | 300 |
| Dose (e-/Å<sup>2</sup>) | <100 | <100 | <100 | <100 | <100 | <100 |
| Defocus (μm) | -2.3 to -3 | -3 | -2.5 to -3.4 | -3 to -5 | -2.5 to -3.5 | -2.5 to -3.5 |
| Pixel Size (Å) | 2.17 | 2.17 | 2.17 | 2.17 | 2.17 | 2.17 |
| Increment (°) | 2 or 3 | 3 | 3 | 2 | 3 | 3 |
| Tilt range* (°) | +/- 54 or 58 | +/- 54 or 57 | +/- 54 | +/-53 | +/- 54 | +/- 54 |
| # of tilt series | 39 | 37 | 21 | 28 | 8 | 28 |
| Cryo-FM | CorrSight | CorrSight | Meteor 50x/0.5<br>NA | Meteor<br>50x/0.8 NA | Meteor 50x/0.8 NA | Meteor 50x/0.8 NA |
| Milling strategy | Manual | Manual | Manual | Manual | Automated | Automated |

### Cryo-electron tomography data acquisition

Lamellae were imaged either on a 200 kV TEM (Talos Arctica, Thermo Fisher Scientific) equipped with a K2 summit direct electron detector and energy filter (Gatan) or a 300 kV TEM (Titan Krios, Thermo Fisher Scientific) equipped with a K3 summit direct electron detector and energy filter (Gatan). Intermediate magnification tilt series were collected at a pixel size of 7.09 Å/pix (Arctica) or 6.32 Å/pix (Krios) and a target defocus of -5 μm. Higher magnification tilt series were collected at a pixel size of 2.17 Å/pix and a target defocus of -3 μm. Tilt series were collected bidirectionally using a dose symmetric scheme, with an increment of 2^0^ or 3^0^ and a total dose of ∼100 e/Å^2^. Generally, tilt series were collected at a tilt range of 54^0^ to -54^0^, while correcting for the lamella angle, which was 9-12^0^, depending on the milling angle. For example, if the lamella was milled at an angle of +9^0^, tilt series were collected from +63^0^ to -45^0^.

### Tomogram reconstruction and template matching

Intermediate magnification tilt series were motion corrected using MotionCor2 (Zheng et al., 2017). Tomogram reconstruction was performed with IMOD v4.10.29 (Kremer et al., 1996). Tilt series were aligned via patch tracking and CTF estimation and correction was performed using the ctfplotter and ctfphase-flip commands. Tomograms were reconstructed at bin4 and denoised using nonlinear anisotropic diffusion (NAD) in IMOD v4.10.29 (Frangakis and Hegerl, 2001). Membrane segmentation was performed on these NAD-denoised tomograms.

For tilt series acquired at high magnification (2.17 Å/pix). The movies of each tilt were motion corrected using MotionCor2 (Zheng et al., 2017) after which the CTF per tilt was estimated in WARP (Tegunov and Cramer, 2019). The pre-processed tilt stacks were aligned and reconstructed in AreTomo (Zheng et al., 2022). For data collected with the Titan Krios with K3 camera, tilt series were cropped into a square shape prior to alignment, as this slightly improved alignment performance. Tomograms were denoised for visualization with CryoCARE v0.3.0 (Buchholz et al., 2019).

### Template matching and subtomogram analysis

To locate the ribosomes within the data, template matching was performed with pytom-match-pick (Chaillet et al., 2023) on 8x binned tomograms (17.36 Å/pix) using an 80S yeast ribosome (emdb: 18231,(Rangan et al., 2024)) as template, and a mask with a radius of 10 pixels (173.6 Å). Template matching results were manually checked in Napari v0.4.17, using the blik v0.6 plugin for data visualization (Gaifas et al., 2024). With the obtained particle coordinates, subtomograms and related CTF volumes were extracted at a pixel size of 8.68 Å/pix (4x binned) in WARP (Tegunov and Cramer, 2019).

Particle coordinates from different sample types and data collections were merged in a single particle list for the entire subtomogram analysis (STA) workflow (Supp. Fig. 3). The 158,222 subtomograms, extracted previously in WARP together with their respective CTF volumes, were aligned in Relion v3.1.4 (Zivanov et al., 2018) against an 80S ribosome reference (emdb 18231, filtered to 40 Å). Next, 3D classification without alignment (T=0.5, N=12, 80S mask and reference) was performed to remove 60S particles and false positives (Total input,Supp. Table 1). The remaining 127,344 particles (Cleaned, Supp. Table 1) were refined in M v1.0.9 using an LSU hard mask at a pixel size of 4.34 Å/pix (Tegunov et al., 2021). After this step, 4x binned subtomograms (8.68 Å/pix) were extracted and aligned in Relion using an LSU mask. These aligned particles were used in parallel for translational state and ER-bound ribosome classification.

### Translational state classification

The 4x binned aligned particles were split into 2 ribosome classes using 3D classification without alignment (T=4, N=2, SSU mask and 80S reference). These two classes (Class 1 and Class 2, Supp. Table 1) were refined separately in M using an LSU hard mask at a pixel size of 2.17 Å /pix. 2x binned subtomograms (4.34 Å/pix) were extracted and aligned in Relion using an LSU mask. To determine the ribosomes’ translational state, 3D classification without alignment and a mask focused on the tRNA binding site was performed (T=4, N=5 for class 1 and N=7 for class 2, own refined 80S reference). Translational states were assigned based on the position of the tRNA’s and elongation factors in PDB and EMDB maps, namely 4UJE (E- and P-site tRNA, (Budkevich et al., 2014)), 4CXG (Sampling tRNA, (Budkevich et al., 2014)), 6Y0G (P- and A-site tRNA, (Bhaskar et al., 2020)), emdb2904 (A- and P/E-site tRNA, (Behrmann et al., 2015)), 6Y57 (A/P- and P/E-site tRNA, (Bhaskar et al., 2020)), 6Z6M (E-site tRNA and eEF2, (Wells et al., 2020)), 6GZ5 (E- and P-site tRNA, and eEF2, (Flis et al., 2018)), 5GAK (eIF5A, (Schmidt et al., 2015)) and 7B7D (eEF3, (Ranjan et al., 2021)). Each classification was repeated 3 times to determine the reproducibility of the translational state assignment. Each state was aligned in Relion using an LSU mask, and a final average was obtained in M using a hard LSU mask at 2.17 Å/pix. Global resolution estimates, for the whole ribosome, were obtained in M, while local resolutions measurements were obtained with Relion.

### ER-bound ribosomes classification

ER-bound particles were classified out from the 4x binned aligned ribosome particles by 3D classification with an ellipsoid mask positioned below the exit tunnel (Supp. Fig. 3C) yielding 12,396 particles (T=4, N=20, 80S reference). The sensitivity and specificity of classifications with 4 different masks were determined (Supp. Fig. 5B-C). The mask resulting in the highest specificity was used for the actual classification (Supp. Fig. 5A-C, #4), see ‘Statistics and data representation’ for the related calculations. Next, the ER-bound ribosomes were aligned using an LSU mask, using the 2x binned subtomograms and alignment parameters that were obtained for these particles during alignment in Relion for the translational state classification. To further improve resolution around the translocon, particles were shifted 5.4 nm up to position the membrane more in the centre of the box. Another round of 3D refinement was then performed using a mask merging the ellipsoid and the LSU mask (Supp. Fig. 3C). A final average was obtained in M using this ellipsoid and LSU mask at 2.17 Å/pix. The global resolution estimate was obtained in M, while local resolutions measurements were obtained with Relion.

### Membrane segmentation and ER volume analysis

ER membranes in tomograms at intermediate magnification were detected using TomoSegMemTV (Martinez-Sanchez et al., 2014). These initial segmentations were manually completed in Avizo v9.2.0 (FEI). ER membranes were distinguished from other organelles mainly based on the attachment of ribosomes. To calculate the luminal ER volume, the segmented volumes were binarized in Avizo and filled using up to 17 iterations of filling and erosion with PyTom v0993 and Scipy functions (Chaillet et al., 2023; Virtanen et al., 2020). To compare cortical ER volumes between different tomograms, we corrected for the available cell area, taking into account volume thickness (∼number of slices) and the total length of plasma membrane (PM) (in the central tomogram slice) (Eq. 1).

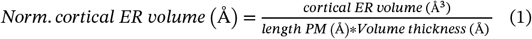

Membrane segmentation for high resolution tomograms was performed on 8x binned CryoCARE denoised tomograms using MemBrainSeg v0.0.2 (Lamm et al., 2024).

### Statistics and data representation

To estimate the performance of the ER-bound ribosome classifications, non-ER-bound ribosomes were manually deselected to generate a ground truth (GT) particle list for 4 tomograms. For each mask, the GT particle coordinates were matched to the particle coordinates of the classified particle lists, yielding a true positive count (TP). The false negative count (FN) was determined by subtracting the TP from the GT. The false positive (FP) count was obtained by subtracting the true positive count from the total number of particles in the classified particle list (total count). The true negative (TN) count was determined by subtracting total cytosolic ribosomes per tomogram from the FN. The TP, FN, FP and TN were used to calculate the sensitivity and specificity of 3D classifications with the different masks (Trevethan, 2017).

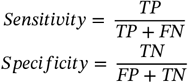

For the calculation of ER volume differences upon ER stress, a Mann Whitney U-test was performed. To compare the abundance of translational states of non-stressed and stressed cells, we performed a Mann Whitney U-test. To compare the translational state abundance of cytosolic ribosomes and ER-bound ribosomes, we performed a Wilcoxon signed-rank test. To compare the differences between the abundance of ribosomes inside and outside vacuoles, a paired T-test was performed. Due to our technical replicates (3 classification runs) each tomogram has 3 different abundance distributions (1 per classification run). These were averaged together so each datapoint had 1 fractional value instead of 3. This way, only the biological replicates were used in the statistical analysis.

To confirm that averaging the technical replicates together in order to perform statistical calculations was appropriate, we also calculated the translational state fractions per classification run. This means that per classification run all particles from the same condition were merged and based on that the translation state fractions were calculated (Supp. Fig. 8). These distributions are comparable to the per-tomogram distributions, and confirmed that indeed the variability between technical replicates is considerably smaller than between biological replicates.

Graphs were created with Jupyter notebook 6.0.3 (Kluyver et al., 2016). Density maps and PDB files were visualized in ChimeraX (Pettersen et al., 2021). Tomograms were visualized in IMOD (Kremer et al., 1996) while segmentations and plotted ribosomes were visualized in ChimeraX using the ArtiaX plugin (Ermel et al., 2022).

Structural prediction for the yeast translocon was done using AlphaFold 3 (https://alphafoldserver.com) (Abramson et al., 2024). Sequences of 5 translocon components were included in the prediction, namely Sec61p (YLR378C), Irc2p (YEL001C), Sss1p (YDR086C), Ysy6 (YBR162W-A) and Sbh1p (YER087C-B).

## Supplemental Figures and tables

**Supplemental Figure 1:**
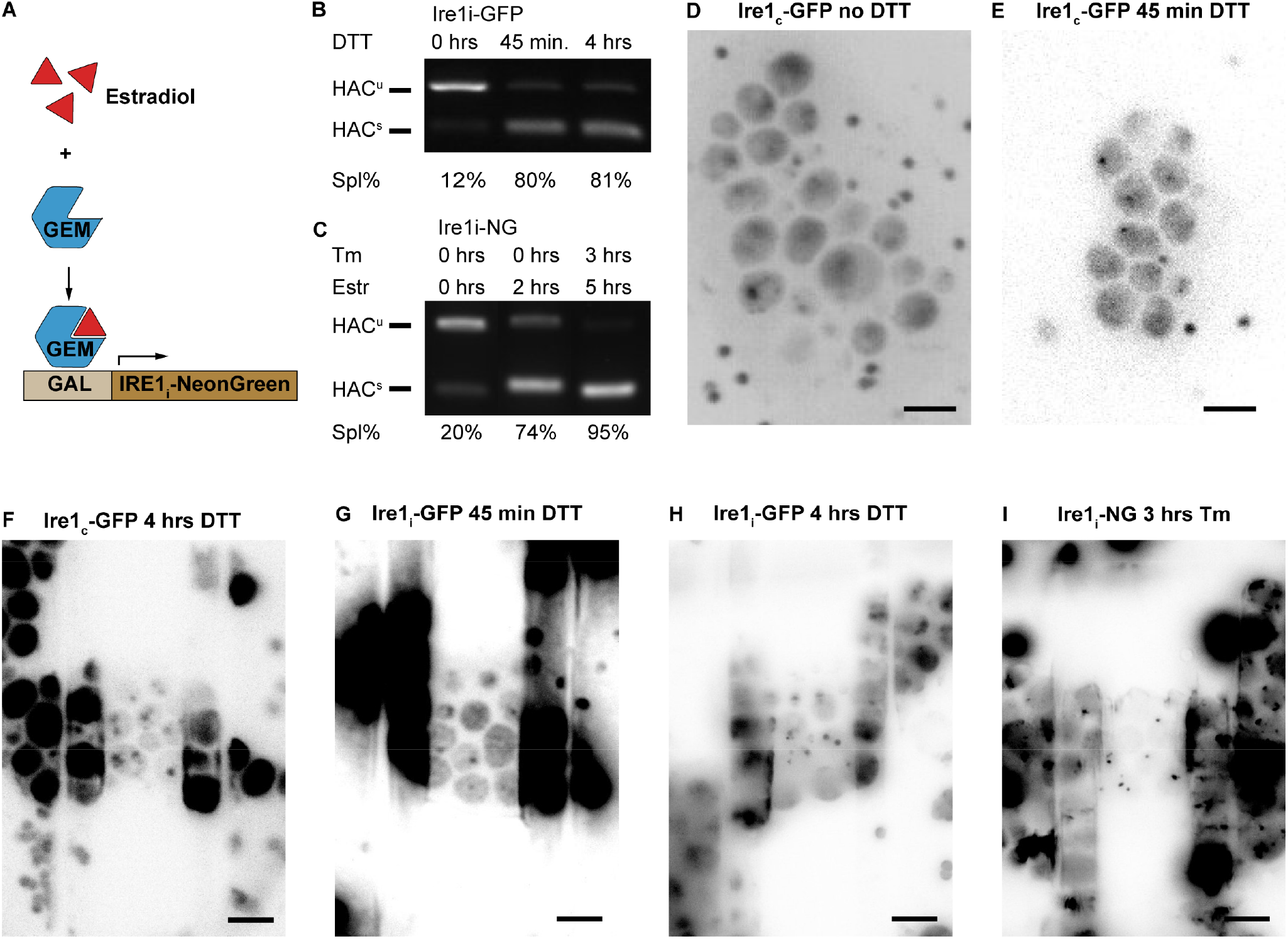
ER stress induction of yeast cell lines. **(A)** Treatment of the Ire1_i_-NG cell line with estradiol leads to activation of the GEM transcription factor. This transcription factor binds the GAL promoter thereby inducing Ire1_i_-NG overexpression. **(B)** RT-PCR of HAC1 mRNA isolated from Ire1_i_-GFP cells treated with DTT (10 mM) for 0 hrs, 45 min. or 4 hrs, or **(C)** from Ire1_i_-NG cells untreated, treated with estradiol (100 nM) for 2 hrs, or with estradiol (100 nM) for 5 hrs and tunicamycin (Tm, 3 µg/mL) for 3 hrs. Unspliced HAC1 (HAC1^u^, 433 bp) and spliced HAC1 (HAC^S^, 188 bp) can be detected. Presence of HAC1^S^ is indicative of ER stress. Splicing percentage is given below the image. **(D)** Cryo-FM data of Ire1_c_-GFP yeast cells on a grid without DTT treatment, **(E)** treated with DTT for 45 min. (CorrSight, intact cells) or **(F)** for 4 hrs (Meteor, milled cells). **(G)** Cryo-FM data of Ire1_i_-GFP yeast cells on a grid treated with DTT for 45 min. or **(H)** 4 hrs (Meteor, milled cells). **(I)** Cryo-FM data for Ire1_i_-NG yeast cells on a grid treated with Tm for 3 hrs (Meteor, milled cells). Scalebar: 5 µm (D-I).

**Supplemental Figure 2:**
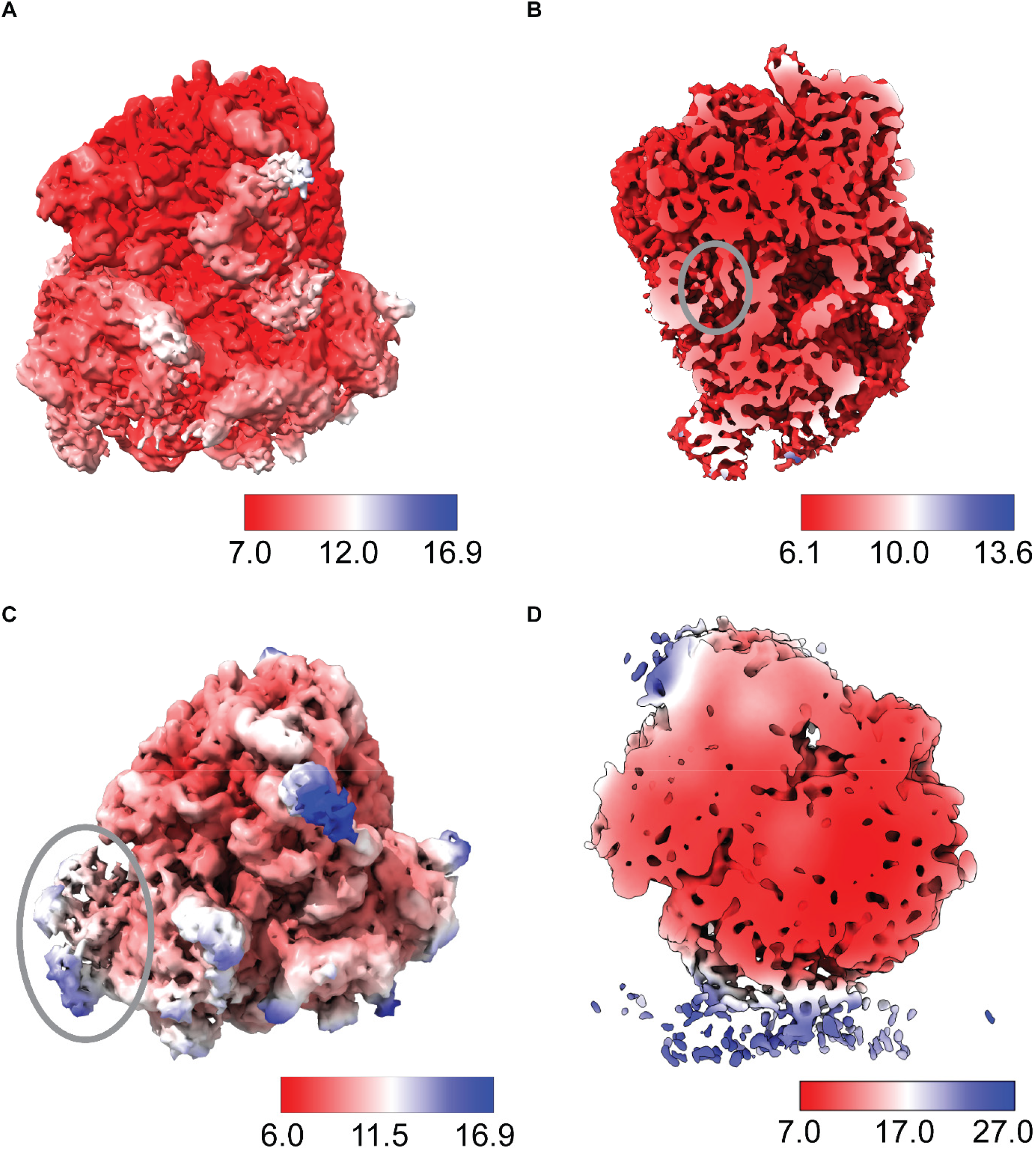
Ribosome local resolution maps. **(A)** Local resolution map of the initial 80S ribosome. **(B)** Local resolution map of the decoding 3 state. The ribosome is clipped to reveal the resolution of the internal parts, including eIF5A (grey circle). **(C)** Local resolution map of the hibernating state, grey circle indicates the location of eEF3. **(D)** Local resolution map of the ER-bound ribosome. The ribosome is clipped to reveal the resolution of the internal parts of the translocon.

**Supplemental Figure 3:**
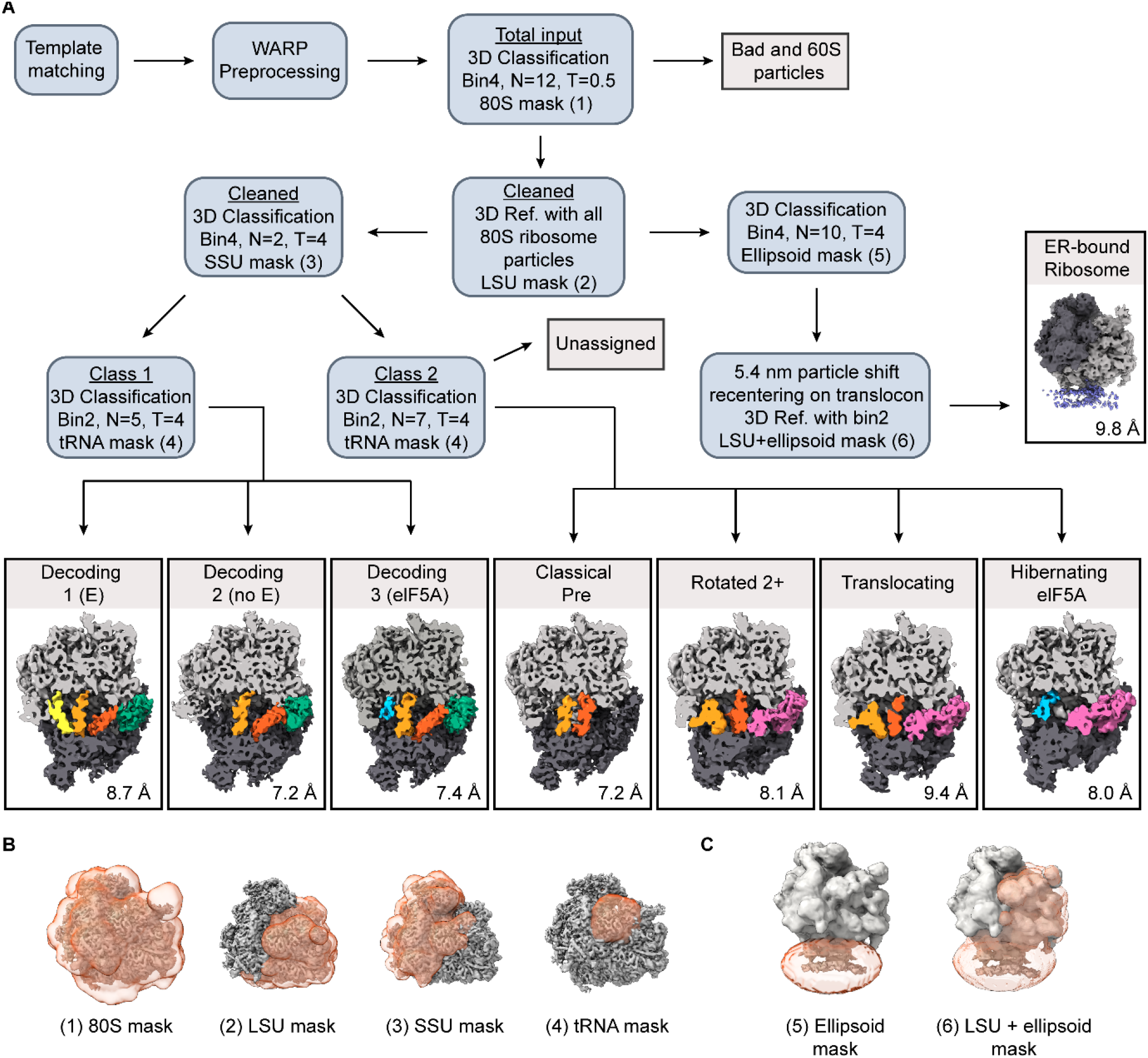
Averaging and classification workflow for translational states and ER-bound ribosomes. **(A)** Diagram describing the different classification and key alignment steps used in this study. Template matching was performed with PyTom. Downstream averaging steps were performed in RELION and M. N = number of classes in RELION classification, T = regularisation parameter. SSU = small subunit, LSU = large subunit. Global resolution, for the whole ribosome, of each state is provided. Underlined terms refer to the used particle list, described in Supp. Table 1. **(B)** Masks used for general alignments and translational state classification. **(C)** Masks used for ER-bound ribosome classification and alignment.

**Supplemental Figure 4:**
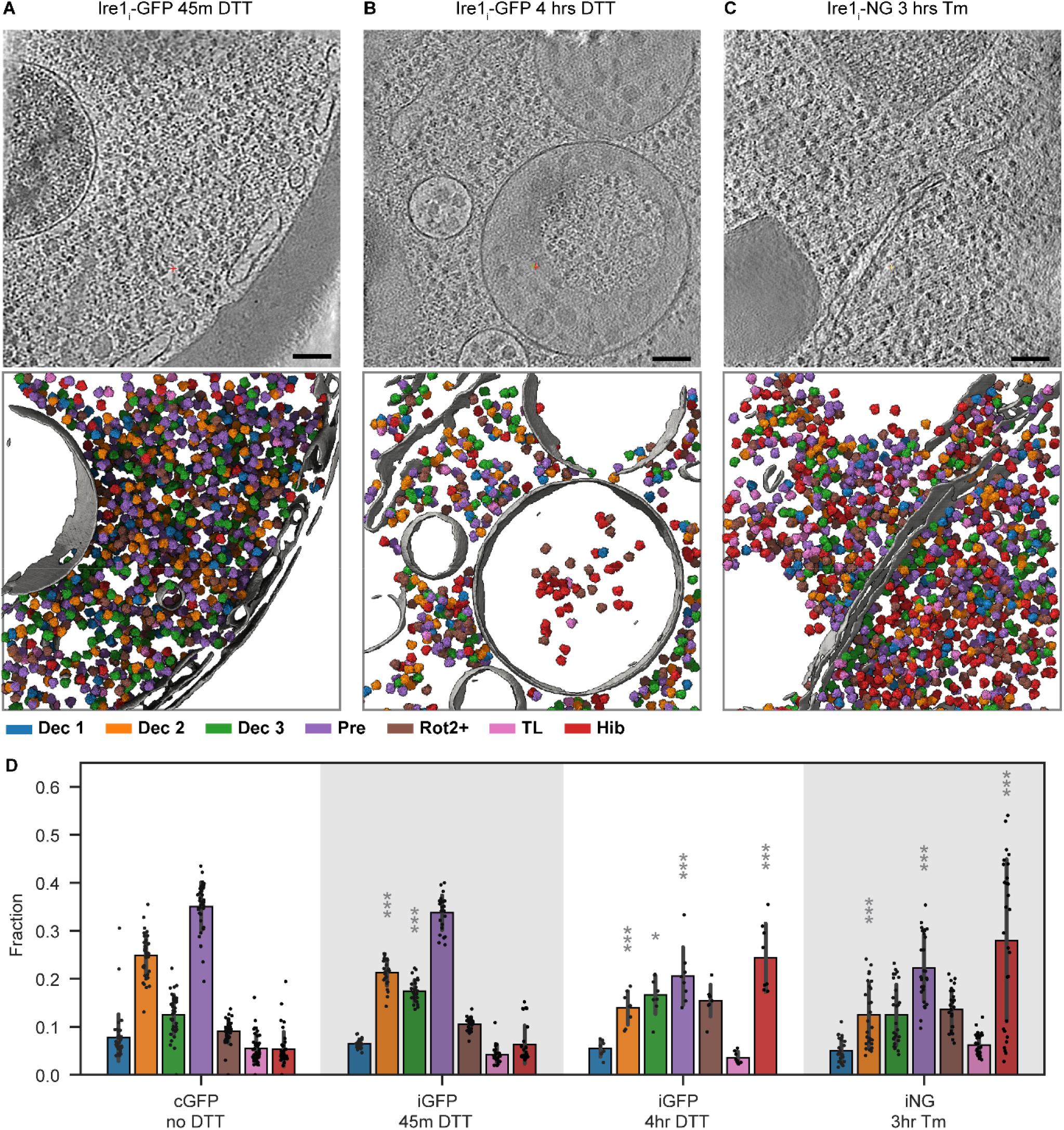
The effect of ER stress on cytosolic translation for 3 additionally stressed samples. Slice through a tomogram of an Ire1_i_-GFP yeast cell stressed with DTT for 45 min. **(A)** or for 4 hrs **(B)**, and an Ire1_i_-NG yeast cell stressed with Tm for 3 hrs **(C)**, including the corresponding membrane segmentation and the mapped-back ribosome particles, which are colour coded according to their translational state. **(D)** The distribution of translational states found in different samples is represented as fractions in a bar plot, cGFP data (Figure 2D) is included as comparison (N=39, 28, 8 and 28 tomograms for no DTT, 45 min. DTT, 4 hrs DTT and 3 hrs Tm respectively). States in stressed samples that had a significantly different abundance in comparison to the non-stressed cGFP sample are indicated with grey asterixis (*P value <0.05, **P value <0.01, ***P value <0.001, Mann Whitney U-test). Scale bar: 100 nm (A-C).

**Supplemental Figure 5:**
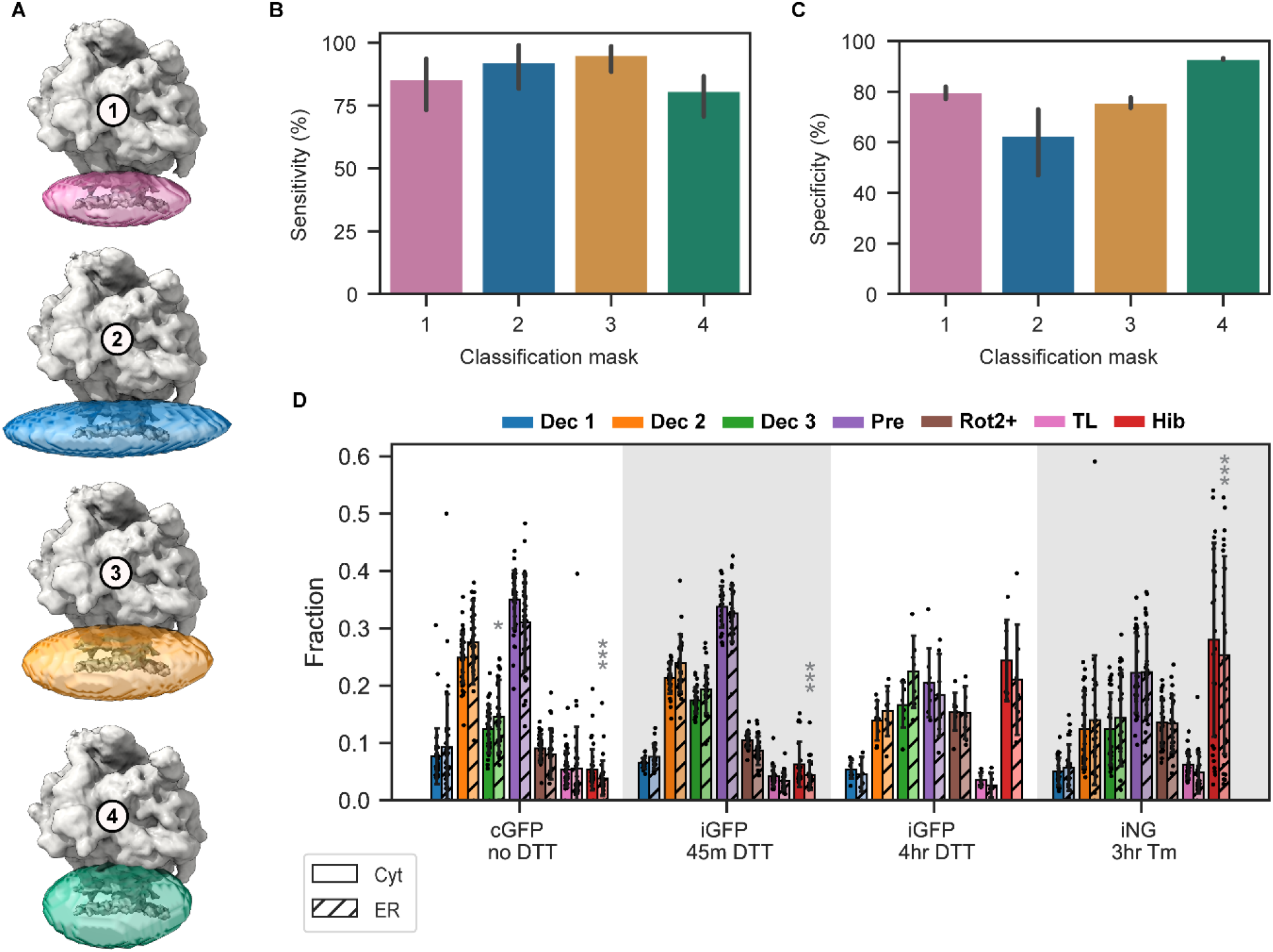
ER-bound ribosome translation under stress in different sample types. **(A)** The 4 different masks that were tested for ER-bound classification. **(B)** Bar plot showing the sensitivity of the 3D classifications using the different masks (N=4 tomograms). **(C)** Bar plot showing the specificity of the 3D classifications using the different masks (N=4 tomograms). **(D)** The distribution of translational states found in different samples is represented as fractions in a bar plot. Both cytosolic ribosome (empty bars) and ER-bound ribosome (hatched bars) fractions are depicted (N=35, 27, 7 and 27 tomograms for no DTT, 45 min. DTT, 4 hrs DTT and 3 hrs Tm respectively). States where the ER-bound abundance is significantly different from the cytosolic abundance are indicated with grey asterixis (*P value <0.05, **P value <0.01, ***P value <0.001, Wilcoxon signed-rank test).

**Supplemental Figure 6:**
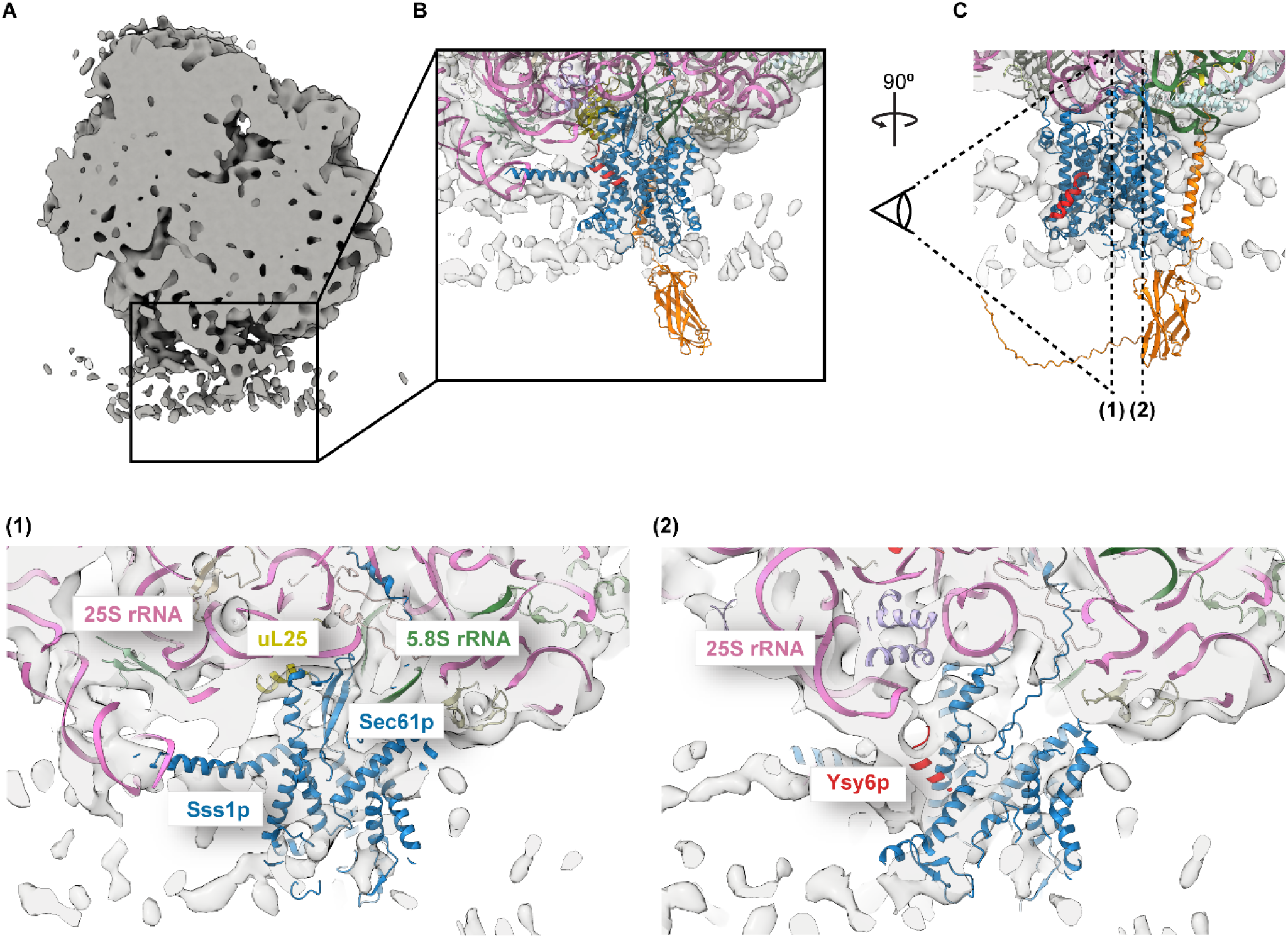
A closer look at the yeast translocon. **(A)** Clipped view of the globally filtered and sharpened ER-bound ribosome map. **(B)** Zoom in on the density beneath the exit tunnel with a ribosome-bound translocon (PDB 3J77 and AlphaFold model, see M&M) fitted into the density. **(C)** Same ribosome with fitted model rotated 90^0^. The dotted lines indicate the point at which the cross section views **(1** and **2)** were made. The relevant translocon and ribosome components are highlighted in different colours.

**Supplemental Figure 7:**
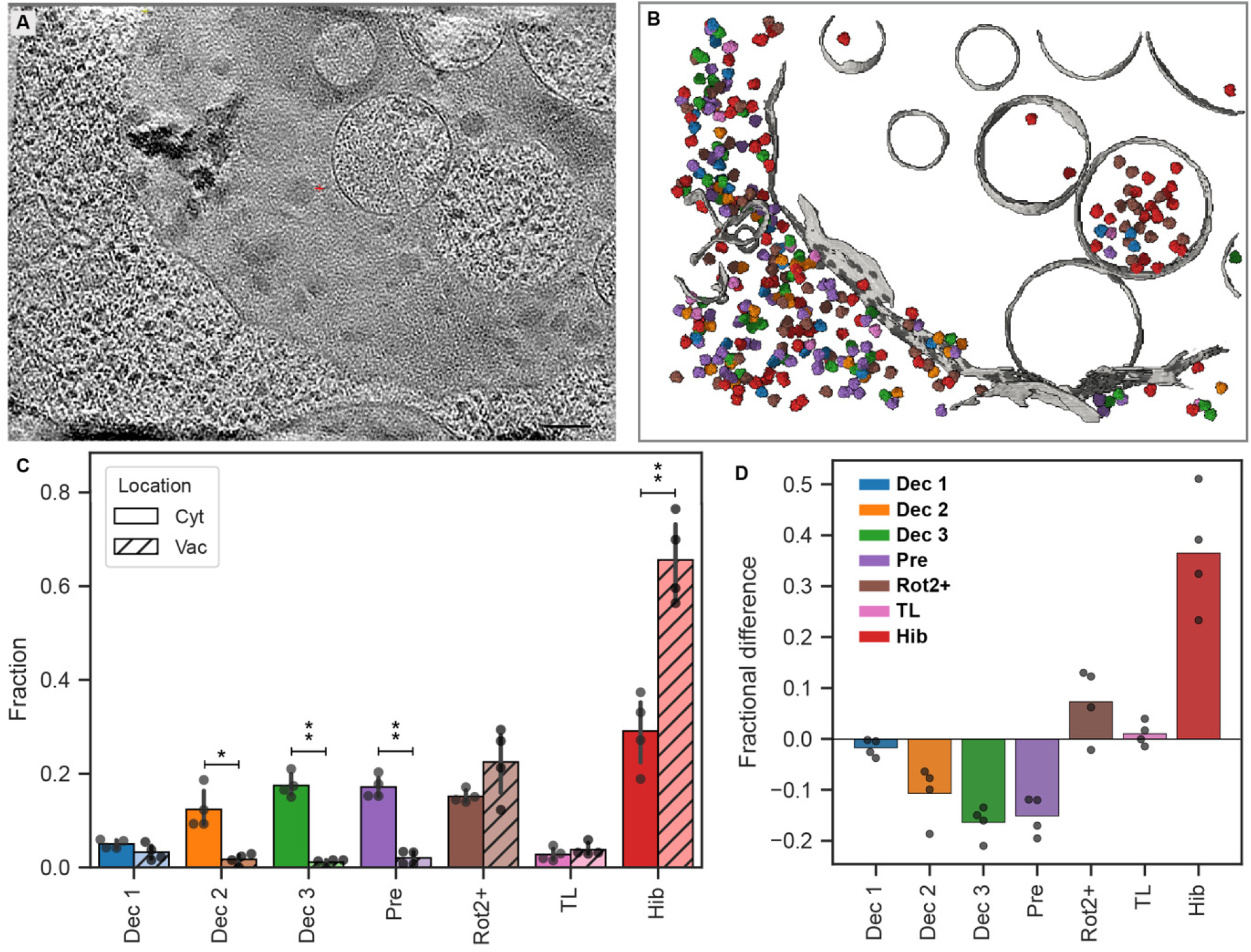
Vacuolar-residing ribosomes in Ire1_i_-GFP cells stressed with DTT for 4 hrs are predominantly hibernating. **(A)** Slice through a tomogram of the cytoplasm of an Ire1_i_-GFP yeast cell stressed with DTT for 4 hrs showing vacuolar-residing ribosomes. **(B)** The corresponding membrane segmentation and the mapped-back ribosomes, which are colour coded according to their translational state. **(C)** Bar graph showing the fraction of each translational state for ribosomes located in the cytosol (empty bars) or vacuole (hatched bars). Individual data points are indicated with a dot (N=4 tomograms). States that have a significantly different abundance between cytosolic and vacuolar ribosomes are indicated with asterixis (*P value <0.05, **P value <0.01, paired T-test). **(D)** Fractional difference between the vacuolar and cytosolic ribosomes, subtracting vacuolar abundance from cytosolic abundance. Scale bar: 100 nm (A).

**Supplemental Figure 8:**
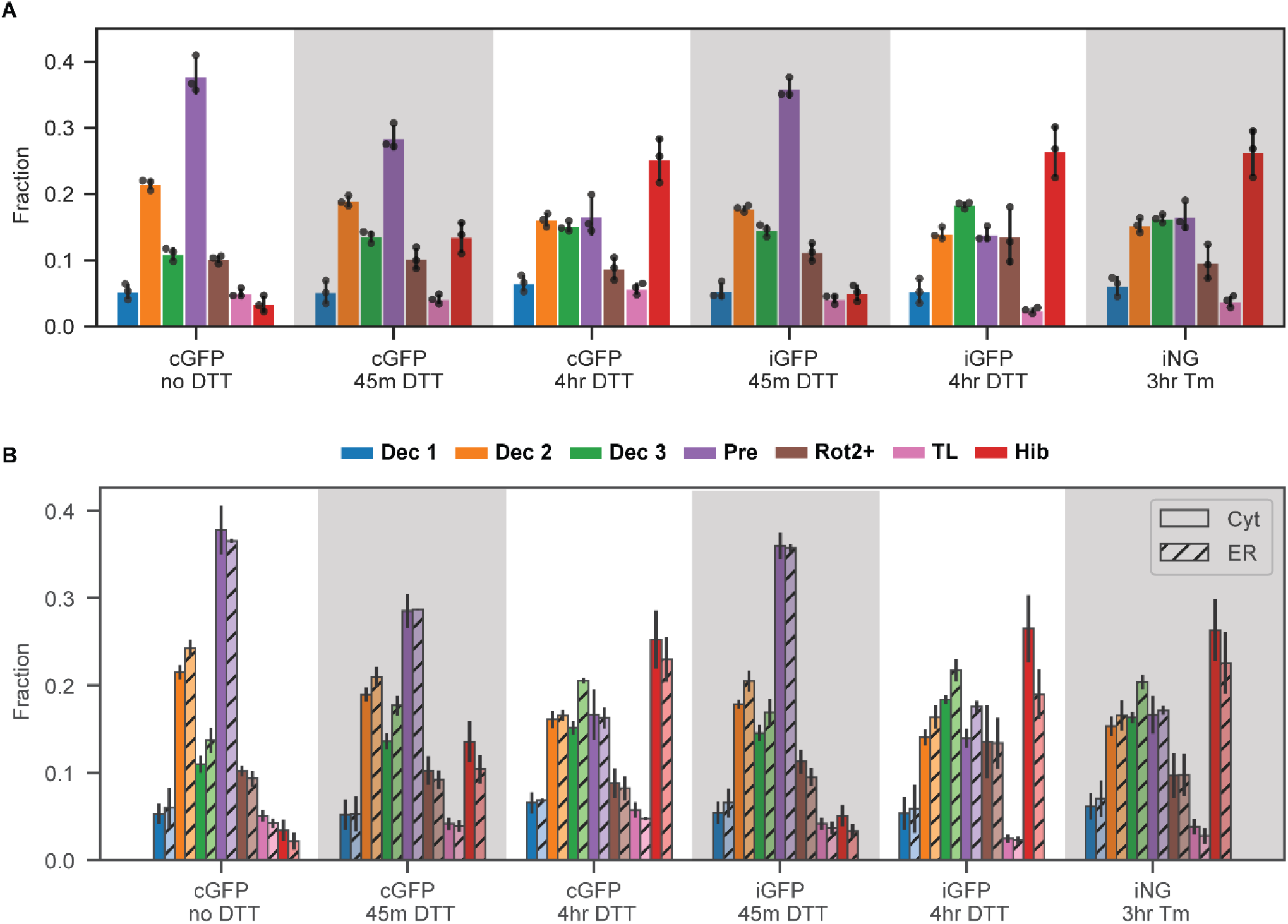
Variation between classification runs (technical repeats). Bar graphs show the distribution of translational states in the cytosol **(A)**, and in both the cytosol and the ER **(B)** per condition. Here, fractions are calculated after merging all ribosomal particles from all tomograms of a condition together, instead of calculating the fractions per tomogram. Data points represent the classification runs (N=3 technical repeats).

**Supplementary table 1:** particle numbers per sample state during the initial processing steps.

|  | cGFP<br>no stress | cGFP<br>45m DTT | cGFP<br>4hrs DTT | iGFP<br>45m DTT | iGFP<br>4hrs DTT | iNG<br>3hrs DTT | Total |
| --- | --- | --- | --- | --- | --- | --- | --- |
| *Total input* | 32,882 | 36,385 | 14,995 | 36,184 | 3,573 | 34,203 | 158,222 |
| *Cleaned* | 22,459 | 28,006 | 11,064 | 34,028 | 3,396 | 28,391 | 127,344 |
| *Class 1* | 10,287 | 11,999 | 4,212 | 16,945 | 1,195 | 8,467 | 53,105 |
| *Class 2* | 12,172 | 16,007 | 6,852 | 17,083 | 2,201 | 19,924 | 74,239 |

**Supplementary table 2:** Total particle numbers per sample state (ER + Cyt) for the different translational states and the 3 different classification repeats.

|  | Repeat | cGFP<br>no stress | cGFP<br>45m DTT | cGFP<br>4hrs DTT | iGFP<br>45m DTT | iGFP<br>4hrs DTT | iNG<br>3hrs DTT | Total |
| --- | --- | --- | --- | --- | --- | --- | --- | --- |
| **Dec 1** |  |  |  |  |  |  |  |  |
|  | 1 | 1,653 | 2,068 | 795 | 2,874 | 222 | 1,547 | 9,159 |
|  | 2 | 1,347 | 1,521 | 682 | 1,962 | 156 | 1,357 | 7,025 |
|  | 3 | 1,008 | 1,034 | 552 | 1,955 | 106 | 927 | 5,582 |
| **Dec 2** |  |  |  |  |  |  |  |  |
|  | 1 | 5,564 | 5,594 | 1,680 | 7,483 | 403 | 3,169 | 23,893 |
|  | 2 | 5,159 | 5,457 | 1,555 | 7,717 | 420 | 2,932 | 23,240 |
|  | 3 | 5,543 | 5,928 | 1,736 | 7,239 | 457 | 3,391 | 24,294 |
| **Dec 3** |  |  |  |  |  |  |  |  |
|  | 1 | 2,518 | 3,827 | 1,547 | 5,685 | 539 | 3,310 | 17,426 |
|  | 2 | 2,982 | 4,356 | 1,676 | 6,242 | 575 | 3,558 | 19,389 |
|  | 3 | 2,895 | 4,217 | 1,567 | 6,498 | 564 | 3,401 | 19,142 |
| **Pre** |  |  |  |  |  |  |  |  |
|  | 1 | 8,473 | 8,391 | 2,316 | 11,121 | 582 | 6,399 | 37,282 |
|  | 2 | 7,322 | 7,645 | 1,750 | 11,008 | 532 | 5,339 | 33,596 |
|  | 3 | 7,016 | 7,543 | 1,657 | 10,913 | 539 | 5,144 | 32,812 |
| **Rot2+** |  |  |  |  |  |  |  |  |
|  | 1 | 2,215 | 3,250 | 1,215 | 3,660 | 676 | 4,194 | 15,210 |
|  | 2 | 2,054 | 2,741 | 994 | 3,457 | 501 | 3,131 | 12,878 |
|  | 3 | 1,847 | 2,404 | 802 | 3,011 | 386 | 2,489 | 10,939 |
| **TL** |  |  |  |  |  |  |  |  |
|  | 1 | 967 | 1,333 | 745 | 1,339 | 106 | 1,536 | 6,026 |
|  | 2 | 910 | 957 | 537 | 1,036 | 79 | 934 | 4,453 |
|  | 3 | 1,124 | 1,121 | 667 | 1,383 | 96 | 1,298 | 5,689 |
| **Hib** |  |  |  |  |  |  |  |  |
|  | 1 | 459 | 2,969 | 2,535 | 1,079 | 827 | 7,509 | 15,378 |
|  | 2 | 626 | 3,762 | 2,868 | 1,568 | 1,014 | 8,849 | 18,687 |
|  | 3 | 893 | 4,231 | 3,202 | 1,845 | 1,146 | 9,926 | 21,243 |

|  |  |  |  |  |  |  |  |  |
| --- | --- | --- | --- | --- | --- | --- | --- | --- |
| *UA* |  |  |  |  |  |  |  |  |
|  | 1 | 610 | 574 | 231 | 787 | 41 | 726 | 2,969 |
|  | 2 | 2,059 | 1,567 | 1,002 | 1,038 | 119 | 2291 | 8,076 |
|  | 3 | 2,133 | 1,528 | 881 | 1,184 | 102 | 1815 | 7,643 |

## Notes

### Competing Interest Statement

The authors have declared no competing interest.

